# KNexPHENIX: A PHENIX-Based Workflow for Improving Cryo-EM and Crystallographic Structural Models

**DOI:** 10.1101/2025.10.02.680106

**Authors:** Suparno Nandi, Graeme L. Conn

## Abstract

New and improved methods for visualizing complex macromolecules in atomic detail continue to expand structural information in the Protein Data Bank but accurately refining atomic models from experimental maps remains a challenge due to efficiency limitations of current refinement approaches. Standard PHENIX refinement can partially address these limitations with its speed and accessibility but often fails to yield the best model compared to more computationally demanding approaches. We therefore developed “KNexPHENIX”, a customized PHENIX-based workflow, to support optimal macromolecular model building. KNexPHENIX can be used to refine macromolecular structures obtained via cryo-electron microscopy (cryo-EM) or X-ray crystallography, regardless of molecular size or composition. KNexPHENIX was evaluated on deposited structures and *de novo* models and consistently produced models with lower MolProbity scores, indicating improved model stereochemistry, compared to default PHENIX, REFMAC Servalcat, REFMAC, or CERES refinement. Importantly, this was accomplished while maintaining model-to-map correlation for cryo-EM datasets and maintaining or reducing the R_free_-R_work_ difference below accepted thresholds for X-ray crystallographic structures, thus limiting overfitting while preserving refinement accuracy. These results establish the KNexPHENIX workflow as a practical, accessible approach for refining both cryo-EM and crystallographic structures, enabling the generation of high-quality models for deposition and guiding further experimental studies.

## INTRODUCTION

Recent advances in structural biology, particularly in single-particle cryo-EM, have driven a rapid increase in the number of structures deposited in the Protein Data Bank (PDB)^1, 2^. Both X-ray crystallography and cryo-EM are now routinely employed to determine the structures of large cellular or viral macromolecular assemblies (kDa to MDa). While obtaining three-dimensional maps for these complexes is technically demanding and time-consuming, an equally significant challenge lies in the need to accurately build and refine atomic models guided by these maps. These structural models require adherence to accepted validation parameter values, including clashscore, Ramachandran outliers, R_work_/ R_free_, and/ or CC_mask_, to be considered reliable by the scientific community^3-5^. Additionally, the task becomes increasingly difficult at resolutions below ∼3 Å, where manual model building is error-prone and labor-intensive^6^. Notably, over 70% of all the cryo-EM structures deposited in the PDB were determined with maps < 3 Å resolution.

To address these issues, several computational methods have been developed such as EM-Refiner^7^, Rosetta-Phenix^8^, High Ambiguity Driven protein-protein Docking (HADDOCK)^9^, and Correlation-Driven Molecular Dynamics (CDMD)^10^. Although effective, some rely on computationally demanding techniques such as Monte Carlo simulation in EM-Refiner^11^ or molecular dynamics (MD) simulations in CDMD^12^. Additionally, the Rosetta-Phenix pipeline performs poorly for large proteins (>1000 residues)^13^, does not refine RNA, and is computationally intensive, while HADDOCK focuses primarily on improvements in clashscore^9^. These limitations restrict the accessibility of such approaches, particularly for researchers without access to high-performance computing resources^14^. In contrast, PHENIX is one of the most widely used software suites for structure refinement due to its broad accessibility, as well as its speed over other methods due to computational efficiency^15-17^. Nevertheless, prior studies have noted that PHENIX does not always generate final models with optimal quality^8, 10, 17^.

We therefore developed KNexPHENIX by integrating multiple PHENIX modules, including ReadySet, PHENIX refinement, and geometry minimization with customized parameters. KNexPHENIX is designed to enhance both existing X-ray and cryo-EM structures, as well as *de novo* models built into maps. Benchmarking shows KNexPHENIX consistently outperforms default refinement available in PHENIX and REFMAC5 for crystal structures^18^, REFMAC Servalcat for cryo-EM structures^19^, and Cryo-EM Re-refinement System (CERES), an online tool for improving deposited cryo-EM structures by PHENIX refinement with customized parameters^20^. KNexPHENIX can improve models at any resolution, but may be particularly beneficial at moderate to low resolution, and thus represents a computationally efficient and accessible alternative for researchers to enhance the quality of their macromolecule structures.

## METHODS

### Overview of KNexPHENIX workflow

The general framework of KNexPHENIX refinement involves five major steps:

1. Addition of hydrogen atoms to the initial model.
2. Refinement using PHENIX, with specific parameters settings.
3. Geometry minimization, with customized parameters.
4. Removal of hydrogen atoms
5. Final refinement in PHENIX, with modified parameters.

For the studies described in this work, PHENIX version 1.21.1-5286 was used throughout, and all calculations were performed using a single processor. Workflows for KNexPHENIX refinement of existing or *de novo* cryo-EM and crystal structures are described in the following sections. Default cryo-EM and crystal structure PHENIX and REFMAC refinement procedures used in this work, as well as additional information on the typical duration for KNexPHENIX refinement, further explanation for the choice of the workflow stages and parameters, outcomes when varying these stages and parameters, two case studies, and a detailed “how-to” guide with screenshots of the PHENIX interface and are provided in Supplementary Information.

### KNexPHENIX Workflow 1: Refinement strategy for deposited cryo-EM models

After adding hydrogen atoms to the model using *phenix*.*ready_set* to ensure proper geometry^21-23^, *phenix*.*real_space_refine* was used for five cycles using local grid search, global minimization, and B-factor refinement. Restraints such as secondary structure and Ramachandran were applied, but not reference model restraints. Additionally, rotamer restraints were added with the sigma value of 0.35, specifically targeting outliers (*rotamers*.*restraints*.*target=outliers*). Both *rotamers*.*tuneup* and *rotamers*.*fit* were assigned to outliers and poormap. For *pdb_interpretation*, Ramachandran restraints for only peptide bonds were added, but peptide planarity constraints were not enforced. The dihedral function type was set to be determined by the sign of periodicity. Next, one cycle of *phenix*.*geometry_minimization* was used with corrections turned on for bond length, bond angle, dihedral angle, chirality, planarity, parallelity, nonbonded distances, rotamer outliers, and secondary structure restraints. Additionally, Ramachandran restraints were applied to the model in *pdb_interpretation*. Hydrogen atoms were then removed from the minimized model using *phenix*.*pdbtools*. The resulting model was subject to a final round of B-factor refinement with the dihedral function type for the dihedral angles set to all harmonic in *pdb_interpretation*. For all the above steps, parameters not specified were set to their default values.

### KNexPHENIX Workflow 2: Refinement strategy for *de novo* modeling into cryo-EM maps

Starting models were selected based on structural similarity to the existing PDB model for the cryo-EM map, reflecting a likely strategy for new structure determination by cryo-EM. Models were stripped of ligands and waters and fit to the map using *fitmap* in Chimera^24^. Next, hydrogen atoms were added, and the modified model was PHENIX refined as described in the previous section, except that a single round of simulated annealing was included within the five refinement cycles, and the dihedral function type was set as all harmonic in *pdb_interpretation*. If post-refinement CC_mask_ is suboptimal, rotamer restraint parameters can be reverted to default values (rotamer restraints enabled, *rotamers*.*fit* set to outliers and poormap, and no specific setting for *rotamers*.*restraints*.*target, rotamers*.*tuneup*, or sigma). Geometry minimization used the same parameters as before, except that five cycles of minimization were used with the dihedral function type set to all harmonic. Next, hydrogen atoms were removed, followed by five cycles of PHENIX refinement using global minimization, B-factor refinement, and reference model restraints based on the starting model. In *pdb_interpretation*, harmonic restraints on starting coordinates were enabled, the dihedral function type was set to all harmonic, and Ramachandran restraints and peptide planarity constraints were applied. All unspecified parameters were left to default. Notably, if the de novo model sequence differs from the available structure, the sequences can be aligned (e.g. using clustalW^25^), variable residues altered manually, and the corrected model optimized by *phenix*.*sculptor*^26^, followed by the subsequent steps of Workflow 2. However, for high quality maps, final model corrections can be made after completing Workflow 2, by re-refining the model using Workflow 1.

### KNexPHENIX Workflow 3: Refinement strategy for previously refined X-ray crystal structures

Hydrogen atoms were added to the model using *phenix*.*ready_set*, and the modified structure was geometry minimized through two cycles, enforcing corrections for bond length, bond angle, dihedral angle, chirality, planarity, parallelity, nonbonded distances, rotamer outliers, and secondary structure restraints. For minimization, harmonic restraints on starting coordinates were enabled in *pdb_interpretation*, Ramachandran restraints and peptide planarity constraints were applied to the model. Post-minimization, hydrogen atoms were removed as described previously, followed by five cycles of PHENIX refinement (*phenix*.*refine*) using real-space, reciprocal-space, occupancy, and B-factor refinement, and included simulated annealing at the second and fourth (penultimate) rounds of refinement. Additionally, refinement cycles were stereochemistry weighted, with secondary structure restraints turned on. In *pdb_interpretation* for PHENIX refinement, harmonic restraints on starting coordinates were enabled, and Ramachandran restraints and peptide planarity constraints were applied. The *dihedral_function_type* was set to all harmonic for both minimization and refinement. Parameters other than those detailed were used with default settings.

### KNexPHENIX Workflow 4: Refinement strategy to improve *de novo* modeling into X-ray crystallographic maps

For model fitting into maps derived from molecular replacement (MR)^27^, an existing structure was chosen from the PDB database based on similarity (RMSD <1.0) to the structure deposited corresponding to that map, followed by removal of water molecules or ligands from the model. PHASER was used to perform MR using the entire model with sequence identity set to 100%. As before, hydrogen atoms were added to the MR model using *phenix*.*ready_set*, followed by *phenix*.*refine* with strategies as described above for refinement of deposited X-ray crystal structures. However, *reference_model*.*enabled* was set to “true” with the starting model used as reference. Parameters in *pdb_interpretation* were the same as above, except that *ramachandran_plot_restraints*.*enabled* was set to “false”. Geometry minimization parameters were also the same, except that harmonic restraints on starting coordinates were not applied. Post minimization, hydrogen atoms were removed from the model, followed by PHENIX refinement as described above. The *dihedral_function_type* was set to all harmonic for both PHENIX refinement and geometry minimization. Settings not otherwise noted remained unchanged from their defaults for each step. If the PHASER predicted model has a different sequence compared to the reference structure, it can be modified after MR for low resolution structures. However, for high resolution structures, it can be altered after completion of Workflow 4 followed by re-refinement using Workflow 3.

## RESULTS

### KNexPHENIX improves the model quality of existing cryo-EM structure

To test the utility of KNexPHENIX for enhancing the model quality of cryo-EM structures previously deposited in the PDB, we chose thirteen models from the database reflecting diversity in the type of macromolecule (proteins and nucleoprotein complexes) and size (kDa to MDa range), and prioritizing structures determined at moderate resolution and with poor model quality indicators (e.g., higher Clashscore and percentage of Ramachandran outliers; **Supplemental Table S1**). To determine improvement, we compared the MolProbity scores^21^ and CC_mask_ values^4^ between the deposited models and those refined using standard PHENIX, Servalcat, or KNexPHENIX Workflow 1 (**Supplemental Fig. S1A**). PHENIX refined structures collectively show slightly higher MolProbity scores and increased CC_mask_ values, indicating poorer model quality, but with slight improvement in model-to-map fit (**Fig. 1A,B and Supplemental Table S2**). Similarly, Servalcat generates the best CC_mask_ values among all the methods tested, but with the poorest MolProbity scores among all re-refined structures. In contrast, KNexPHENIX significantly lowered MolProbity scores while generally maintaining the model-to-map fit in a range similar to the deposited structures, thereby achieving a compromise between the model quality and map fit (**Fig. 1A,B**).

**Fig. 1.**
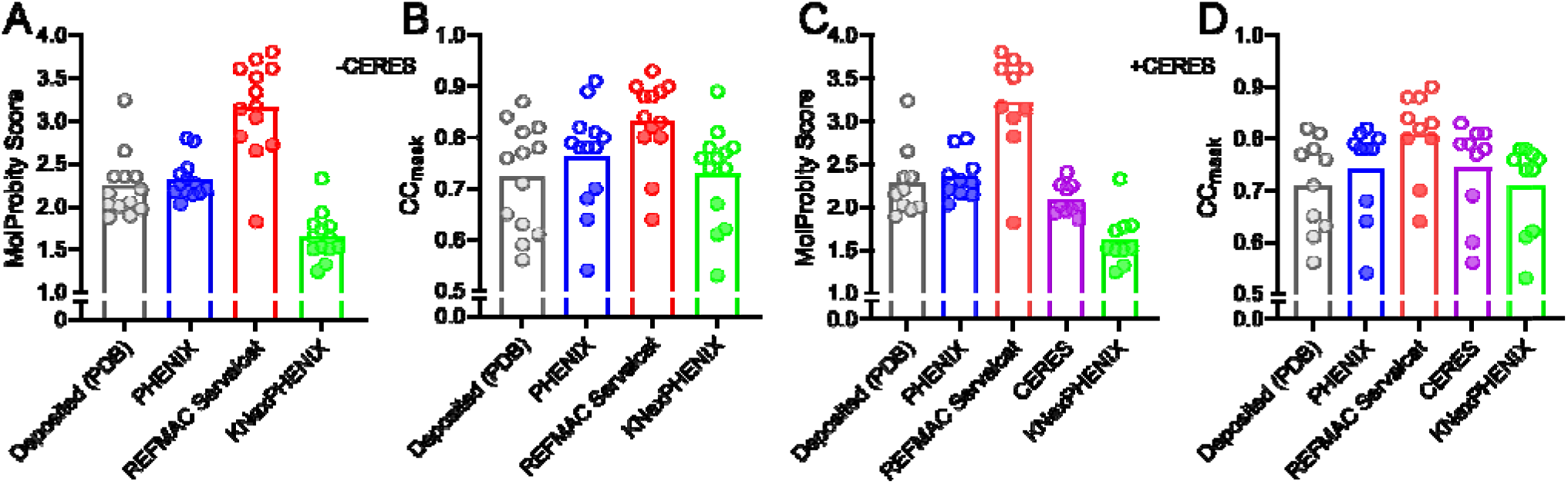
KNexPHENIX enhances deposited cryo-EM model quality without affecting model-to-map fit. ***A***, KNexPHENIX, but not PHENIX or Servalcat, reduces the MolProbity score of the 13 re-refined structures compared to the deposited PDB models. ***B***, CC_mask_ remains consistent with KNexPHENIX and show modest improvement for PHENIX and Servalcat compared to the PDB structures. ***C***, CERES improves the MolProbity score of the 10 test structures compared to PHENIX and Servalcat, though KNexPHENIX performs best. ***D***, The CC_mask_ values for PHENIX, CERES and KNexPHENIX remain consistent while Servalcat shows improvement.

Among the thirteen structures selected for our analyses, 10 were available on the CERES platform, and we therefore additionally compared MolProbity scores and CC_mask_ values on CERES to those from refinement in PHENIX, Servalcat, and KNexPHENIX. Although CERES produces improved model quality based on MolProbity scores compared to PHENIX and Servalcat, KNexPHENIX is capable of further improvement (**Fig. 1C,D and Supplemental Table S3**). However, the model-to-map fit for CERES-refined structures is better compared to KNexPHENIX, with Servalcat performing best in this metric (**Fig. 1C,D**). Overall, these analyses indicate that KNexPHENIX generates models that fit their corresponding maps at least comparably to the deposited model, but with improved MolProbity scores compared to PHENIX, Servalcat, or CERES.

### KNexPHENIX refinement of initial cryo-EM models enhances final model quality

To determine whether KNexPHENIX improves initial cryo-EM models to produce a final structure suitable for PDB deposition, we tested KNexPHENIX Workflow 2 (**Supplemental Fig.S1B**) on ten structures and compared the results with those obtained from PHENIX and Servalcat. This structure set included nine of the thirteen models analyzed previously and one new model that we have recently published^28^ using the method described here, but which underwent additional manual model building post-refinement (**Supplemental Table S1**). The selection was again based on ensuring diversity of macromolecule type and size. The initial models for fitting and refinement to the deposited maps were chosen based on their structural similarity to the corresponding deposited structures. Again, KNexPHENIX consistently genarates models with lower MolProbity scores compared to PHENIX and Servalcat, while Servalcat achieves the best model-to-map fit (**Fig. 2A,B and Supplemental Table S4**). However, the CC_mask_ for both PHENIX and KNexPHENIX is similar (**Fig. 2B**). Where additional manual adjustment of the KNexPHENIX-refined model is necessary, refinement may be continued using the workflow applied for deposited structures. We also tested KNexPHENIX with alternative starting models using two predicted structures from AlphaFold (AF)^29^, which again showed improvement over the PDB models but exhibited minor differences. Therefore, we extended this comparison using models for PDB 8GUD from the Boltz2^30^ and RoseTTAFold3 (RF3)^31^ servers (predictions for PDB 5A1A were not possible as it is >3,500 residues). These analyses produced similar results to AF, indicating the suitability of integrating AF, Bolt2, or RF3 predictions with KNexPHENIX (**Supplemental Table S5**). Overall, KNexPHENIX generates improved model quality with negligible effect on map-to-model fit compared to standard PHENIX refinement.

**Fig. 2.**
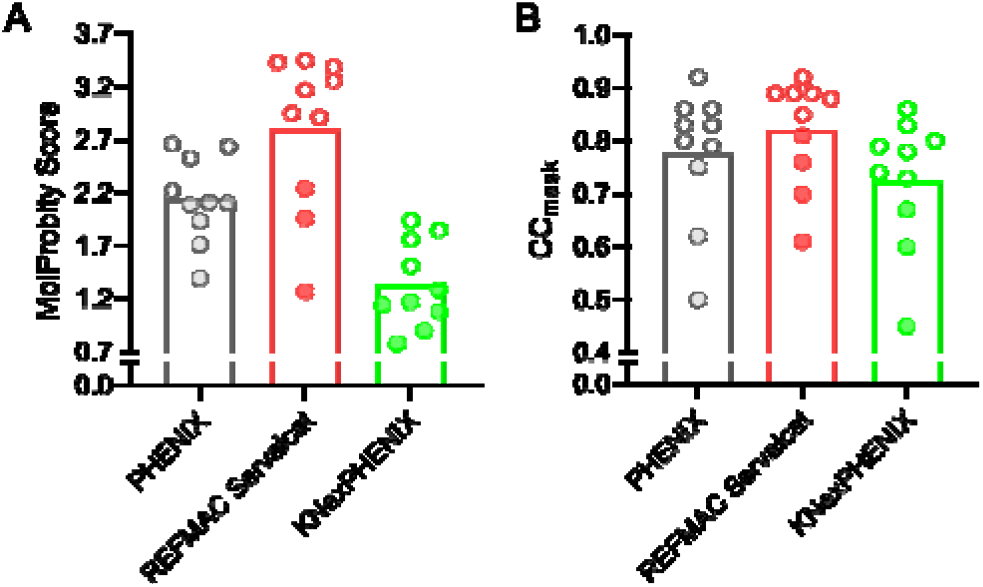
Refinement of initial cryo-EM models using KNexPHENIX enhances model quality. ***A***, MolProbity scores of the 10 structures refined by KNexPHENIX are significantly better than the initial models processed by PHENIX or Servalcat. ***B***, CC_mask_ from Servalcat is higher than the other two refinement procedures.

### Improving model quality of deposited X-ray crystal structures using KNexPHENIX

Next, to assess whether KNexPHENIX can improve existing crystallographic models in the PDB, we tested its performance on sixteen protein structures using KNexPHENIX Workflow 3 (**Supplemental Fig. S2A**). Test structures were again chosen based on their type (proteins and protein-protein complexes), size (∼15 to ∼230 kDa), and displaying indicators of suboptimal model quality and resolution (**Supplemental Table S1**). Along with MolProbity score, values for R_work_, R_free_, and R_free_-R_work_, which can indicate model overfitting^32, 33^, were used to determine the effectiveness of KNexPHENIX in comparison to the deposited structures as well as re-refinement using PHENIX and REFMAC5.

As for cryo-EM structures, KNexPHENIX improves existing models and performs better than PHENIX and REFMAC5 in terms of MolProbity score (**Fig. 3A and Supplemental Table S6**). Both PHENIXand REFMAC5 lower the R_work_ for the existing models, while this metric is unchanged for KNexPHENIX (**Fig. 3B**); in contrast, R_free_ values of the deposited structures are comparable for both REFMAC5 and KNexPHENIX and slightly elevated by PHENIX (**Fig. 3C**). Importantly, KNexPHENIX refinement consistently restricts the difference between free and work R-factors (R_free_-R_work_) to below 5, widely considered an acceptable value^33, 34^. In contrast, due to their greater impact on R_work_ compared to R_free_, the average R-factor difference for the PHENIX and REFMAC5 refined models is closer to 6 (**Fig. 3D**). Altogether, our results show that KNexPHENIX can enhance the quality of the existing structures obtained by X-ray crystallography, while limiting overfitting of the model into the map and maintaining model correctness.

**Fig. 3.**
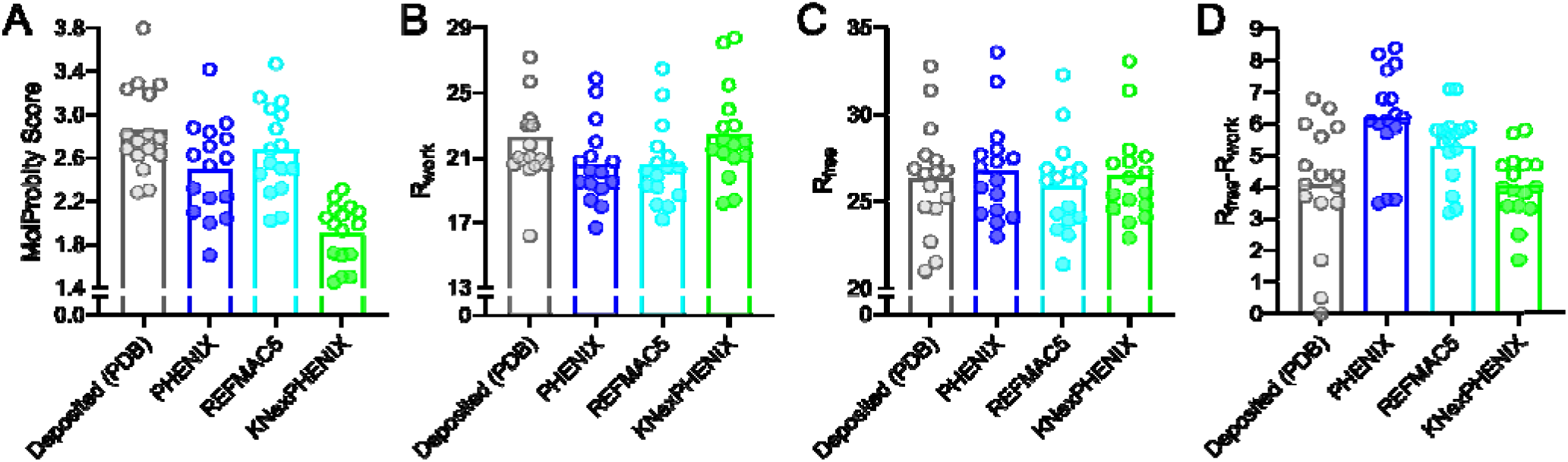
KNexPHENIX enhances existing crystal structure quality while maintaining the map-to-model fit. ***A***, KNexPHENIX significantly lowers the MolProbity score compared to PHENIX, REFMAC5, and the 16 deposited structures. ***B***, Both PHENIX and REFMAC5 exhibit lower R_work_ compared to KNexPHENIX and deposited structures ***C***, R_free_ stays consistent for REFMAC5 and KNexPHENIX, while PHENIX increases this metric. ***D***, PHENIX and REFMAC5 increase the difference between R_work_ and R_free_, whereas it is unaltered by KNexPHENIX.

### MR models for crystallographic maps can be improved by KNexPHENIX

To assess whether KNexPHENIX improves final models in MR-based structure determination, we selected ten structures from the set of sixteen analyzed above. These test cases were chosen based on good initial model-to-map fit and similarity to the deposited models of the MR model. As with other refinment methods, KNexPHENIX could not produce a better final model if this condition was not met, indicating that an adequate initial fit is essential for successful refinement.

KNexPHENIX refinement of these initial models resulted in a final model with improved MolProbity score compared to PHENIX and REFMAC5 (**Fig. 4A and Supplemental Table S7**). Again, the lowest R_work_ was obtained with these two methods, and REFMAC5 produced models with the lowest R_free_, while KNexPHENIX produced models with the lowest difference between R_free_ and R_work_ (**Fig. 4B-D**). Unlike successful refinement of predicted cryo-EM structures, KNexPHENIX refinement of selected crystal structures using starting models from AF, Boltz2, or RF3 resulted in greater variability of validation parameters compared to the refinement of the chosen initial models for the corresponding structures, highlighting the need to sample different initial models prior to refinement (**Supplemental Table S5**). In cases where the user aims to perform further manual corrections, the model obtained from KNexPHENIX can subsequently be processed and further improved using the strategy outlined above for deposited crystal structures. In general, KNexPHENIX generates final models with better quality compared to other methods, effectively preventing overfitting to the map and maintaining model accuracy.

**Fig. 4.**
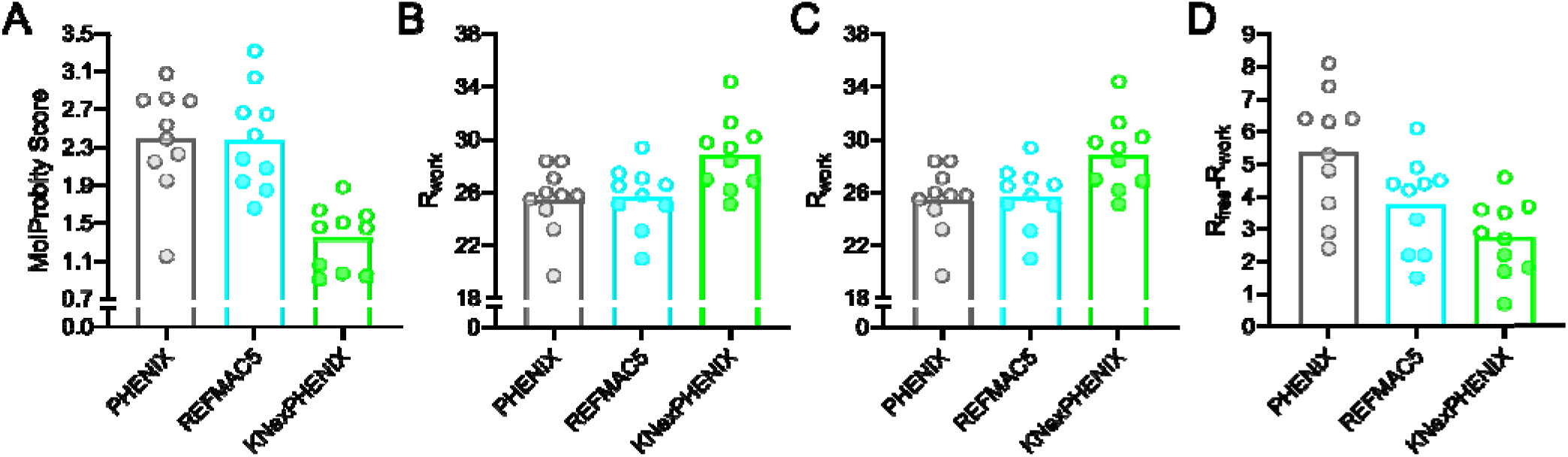
KNexPHENIX refinement of MR models improves X-ray crystallographic structure quality. ***A***, KNexPHENIX improves the *de novo* model quality of 10 models compared to REFMAC5 or PHENIX. ***B***, R_work_ is slightly decreased with PHENIX and REFMAC5. ***C***, R_free_ is lower for REFMAC5 compared to the other two methods. ***D***, R_free_-R_work_ is lower in KNexPHENIX than the other methods.

## DISCUSSION

Efficient and accurate model building remains a key challenge in structural biology, especially for large molecules and/ or poor-quality maps. Although several high-performance computing tools exist, many are computationally intensive and inaccessible to researchers with limited resources. To address this, we developed KNexPHENIX, a set of PHENIX-based workflows that use existing programs of the PHENIX suite with customized parameters to rapidly and efficiently improve the quality of X-ray crystallographic and cryo-EM structural models.

Visual analysis of the H1047R variant of PI3Kalpha (PDB code 8GUB) demonstrates that KNexPHENIX subtly adjusts side chain conformations to improve geometry without altering the model-to-map fit (**Supplemental Figs. S3A and S4A**). In contrast, REFMAC Servalcat produced minimal side chain corrections and reduced geometry quality, as reflected by the increased MolProbity score (**Supplemental Fig. S3A**). Although the standard PHENIX refinement adjusted the side chains more extensively, these changes did not noticeably impact the MolProbity score, underscoring the potential of KNexPHENIX as an alternative to the default PHENIX approach (**Supplemental Fig. S3A**).

Similar results were observed in the refinement of the crystal structure of monoubiquitinated PCNA (PDB code 3L0W; **Supplemental Fig. S3B and S4B**) and other test cases. Again, KNexPHENIX improved the MolProbity score through localized adjustments in protein side chain positions. For instance, in the R220A metBJFIXL HEME domain crystal structure, KNexPHENIX shifts the R226 side chain away from T230, increasing the distance between R226 C□ and T230 OH from 2.8 to 3.1 Å, thereby reducing the clash flagged by Probe in the original deposited PDB model (**Supplemental Fig. S3C**). Along with translational movement, side chain rotation also plays a role in reducing the MolProbity score as observed in the repositioning of the M156 side chain away from T250 (**Supplemental Fig. S3D**). Importantly, these side chain movements, as well as the overall model, remain visibly well fit within the map (**Supplemental Fig. S4C,D**), as also reflected by the R_work_ and R_free_ values.

Further inspection of the monoubiquitinated PCNA structure highlights how KNexPHENIX improves the MolProbity score. In the PDB model, Probe identified a clash between one of the methyl groups of L88 and the backbone N atom of T89 (**Supplemental Fig. S3E**). While PHENIX and REFMAC5 fail to resolve this clash by moving the atoms by 3.1 Å and 3.0 Å apart, respectively, KNexPHENIX further increased this separation (to 3.3 Å), effectively resolving the clash while maintaining a good fit to the density (**Supplemental Fig. S3E and Fig. S4E**).

The inclusion of restraints in the workflow improves the MolProbity scores, while refinement strategies maximize the map-to-model fit, ensuring a balanced optimization of model stereochemistry and agreement to experimental map. However, visual inspection of KNexPHENIX refined models prior to deposition is essential to ensure the absence of disagreements with the map. For instance, the rotamer outliers for R206 in the metBJFIXL variant and for L58 in the KDEL receptor variant are expected as they agree with the map. Although KNexPHENIX removes these outliers, the sidechains significantly deviate from the density (**Supplemental Fig. S5**). Given the vital role of geometric constraints in KNexPHENIX, future efforts combining Amber/AFITT^35, 36^, may further enhance the effectiveness of the refinement protocol.

Finally, although resolution is known to influence MolProbity scores, all our refinements were conducted at the resolution reported in PDB, thereby negating the effect of resolution. Additionally, KNexPHENIX genuinely improves the models as shown by their lower clashscores compared to the PHENIX refined structures (**Supplemental Table S8 and S9**).

## CONCLUSION

KNexPHENIX is a compilation of certain PHENIX-based procedures with defined parameters that offer distinct advantages over default PHENIX and REFMAC refinements, particularly in optimizing local geometry without compromising the fit to X-ray or cryo-EM map. This model improvement process is faster compared to manual model corrections, thereby rapidly enhancing structure interpretability and reliability. These enhancements are particularly valuable when accurate side chain placement is crucial for understanding macromolecular mechanisms or guiding structure-based design of small-molecule ligands and inhibitors.

## ASSOCIATED CONTENT

### Data and Software Availability statement

All input models, maps, and software–PHENIX (https://phenix-online.org/download), REFMAC (https://www2.mrc-lmb.cam.ac.uk/groups/murshudov/), Servalcat (https://github.com/keitaroyam/servalcat), CERES (https://cci.lbl.gov/ceres)–are publicly available; PDB accession codes are listed in **Supplemental Table S1**. KNexPHENIX uses tools within the PHENIX suite.

### Supporting Information

Supplementary methods, KNexPHENIX workflow (**Supplemental Fig. S1 and S2**), model improvement examples (**Supplemental Figs. S3-S5**), PDB entries tested (**Supplemental Table S1**), test results (**Supplemental Tables S2-S9**), supplementary results, case studies 1 & 2, and the “how-to” guide are included in the Supplementary Information.

## AUTHOR INFORMATION

### Author contributions

S.N.–conceptualization, methodology, investigation, formal analysis, validation, data curation, visualization, writing-original draft and writing-review and editing; G.L.C.–resources, supervision, funding acquisition, project administration, data curation, visualization, writing-review and editing.

### Notes

The authors declare no competing financial interest.

## ACKNOWLEDGEMENTS

This work was supported by awards R01-GM130135 from the National Institute of General Medical Sciences and R01-AI088025 from the National Institute of Allergy and Infectious Diseases.

## Notes

### Competing Interest Statement

The authors have declared no competing interest.

### Summary of Updates

Several sections has been updated to address reviewer concerns and new information has been added to the supplementary file.

