## Supplementary Information for "KNexPHENIX: A PHENIX-Based Workflow for Improving Cryo-EM and Crystallographic Structural Models"

**This file contains:**

Supplementary Figures S1-S5

Supplementary Tables S1-S9

Supplementary Methods

Default PHENIX refinement for cryo-EM and X-ray crystal structures

Model refinement in REFMAC

Supplementary Results

Motivation for stage and parameter selection in the KNexPHENIX refinement pipeline

Typical duration and efficiency of KNexPHENIX refinement

Analyses of the effect of variation of stages and parameters in KNexPHENIX

Supplementary References

Case Study 1: PI3Kalpha H1047R cryo-EM structure (PDB code 8GUB)

Case study 2: Monoubiquitinated PCNA X-ray crystal structure (PDB code 3L0W)

KNexPHEX “How-to” guide (**Workflows 1-4** with PHENIX screenshots)

**Supplementary Figures**

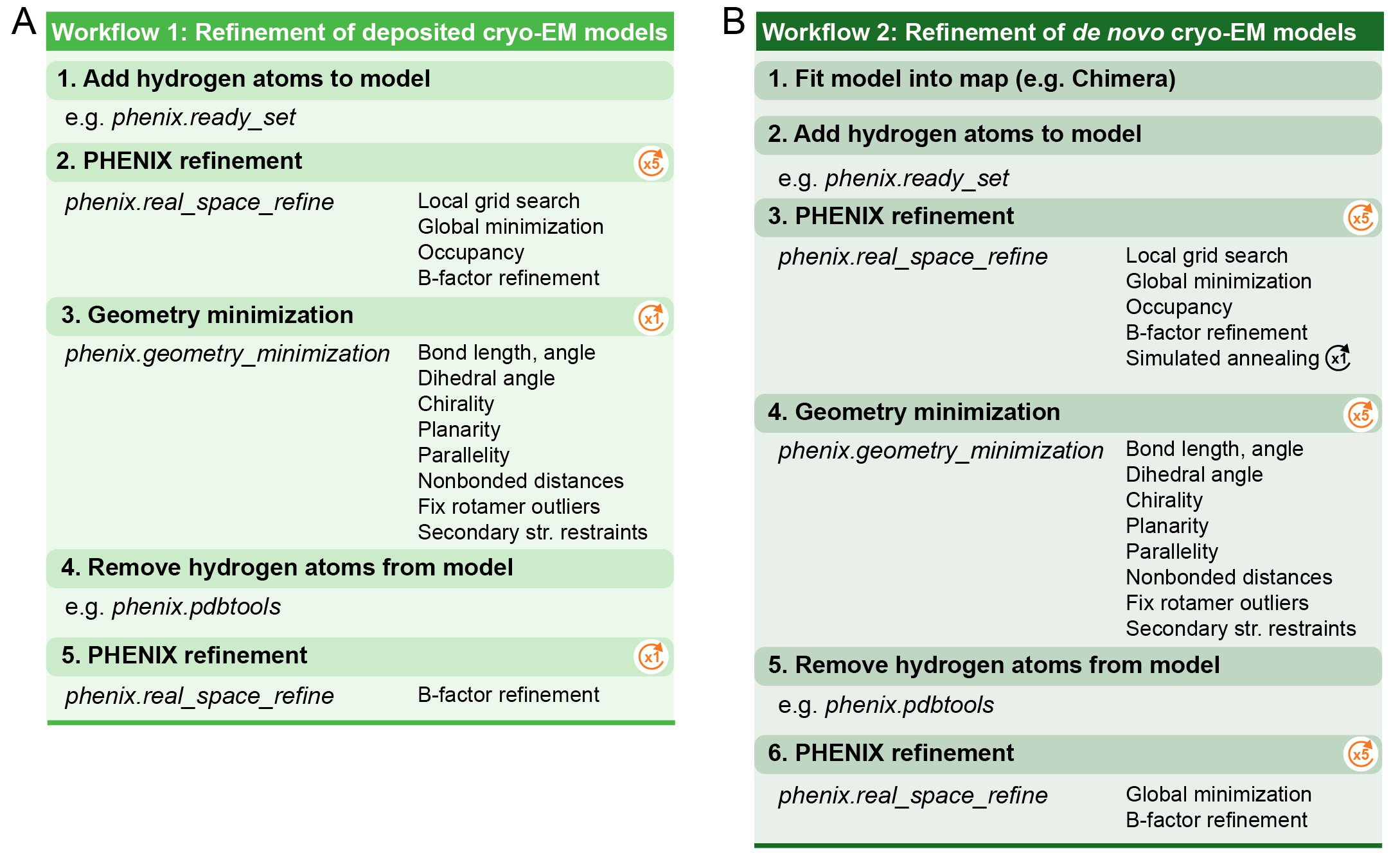

**Fig. S1. KNexPHENIX workflows for refinement of structures determined by cryoEM.** ***A****,* Workflow 1 for refinement of existing cryo-EM models against their corresponding maps. ***B****,* Workflow 2 for refinement of new cryo-EM models in the process of *de novo* structure determination.

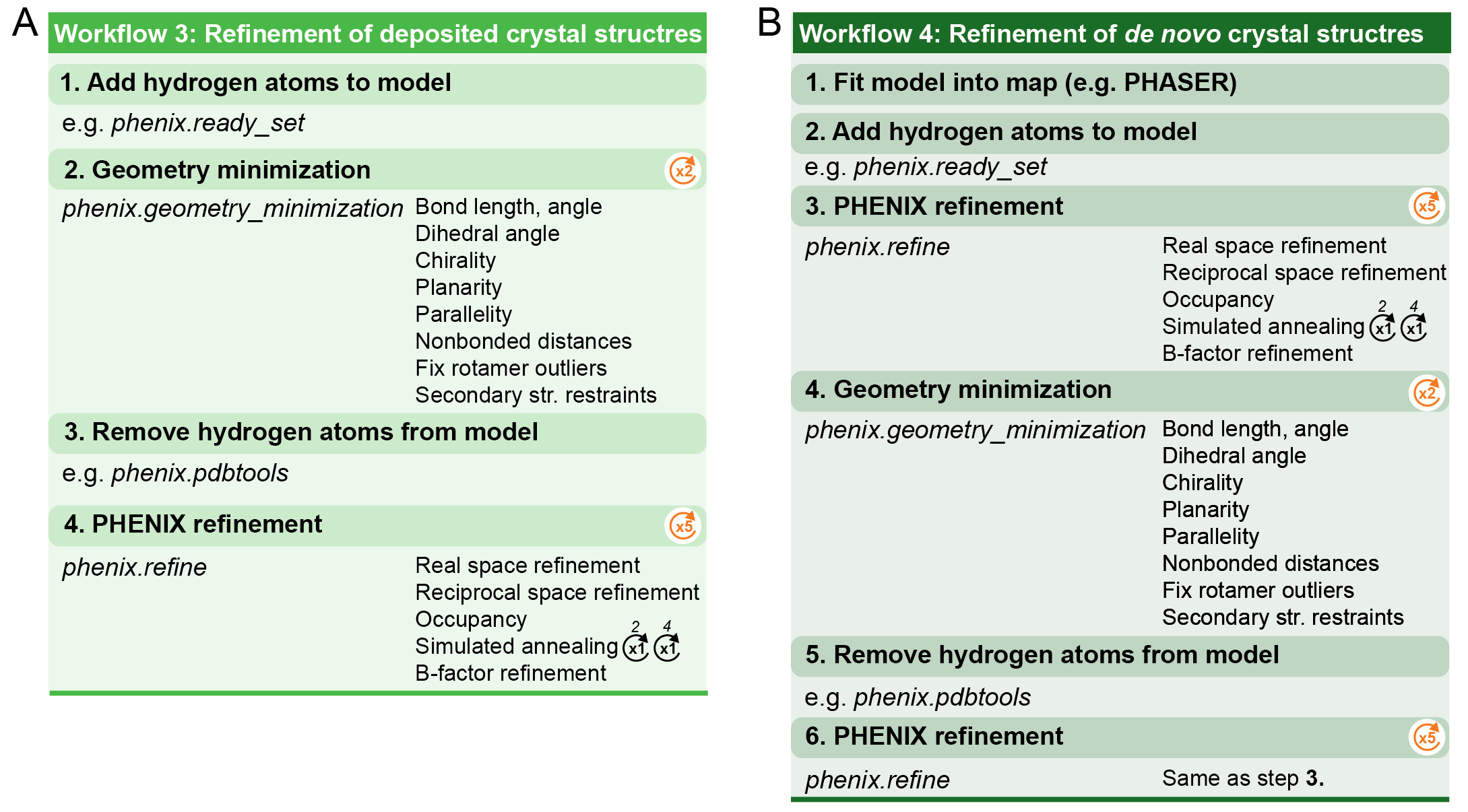

**Fig. S2. KNexPHENIX workflows for refinement of structures determined by X-ray crystallography.** ***A****,* Workflow 3 for refinement of existing X-ray crystal structures against their corresponding maps. ***B****,* Workflow 4 for refinement of new X-ray crystal structures in the process of structure determination by MR.

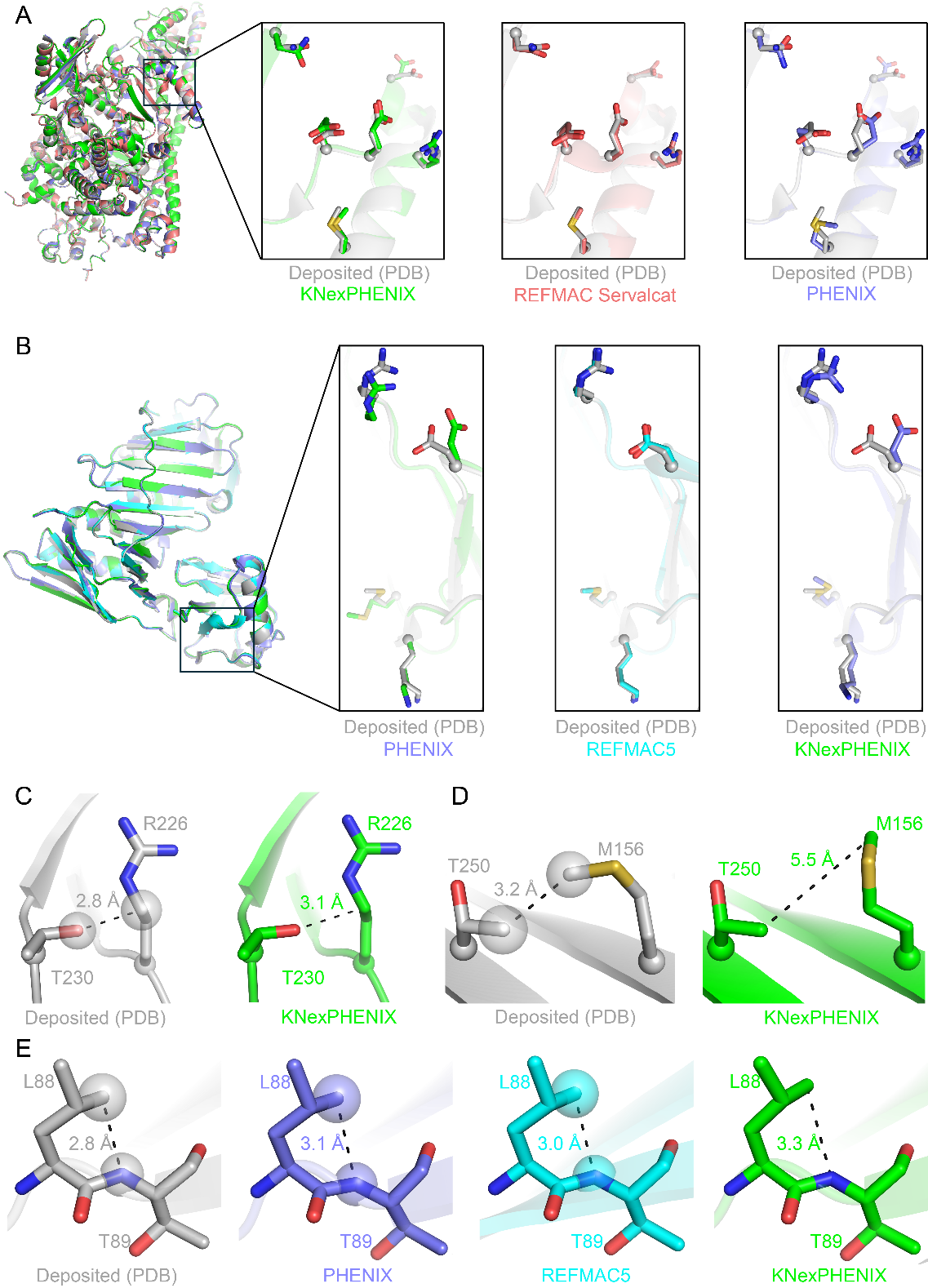

**Fig. S3. KNexPHENIX refines local side chain geometry and resolves atomic clashes without compromising fit to the map. *A,*** Protein backbone conformation is consistent across the structures refined using KNexPHENIX (green), REFMAC Servalcat (red), PHENIX (blue) for the PI3Kalpha H1047R variant determined by cryo-EM (PDB code 8GUB) and all align well with the original PDB model (gray). Sidechain position differences compared to the deposited structure are observed in the structures refined with KNexPHENIX and PHENIX, whereas REFMAC Servalcat leaves them essentially unchanged. ***B,*** The monoubiquitinated PCNA crystal structure (PDB code 3L0W) refined by KNexPHENIX (green), REFMAC5 (cyan), and PHENIX (blue) shows overall structural similarity with the PDB model (gray). REFMAC5 does not change the position of the side chains, while both PHENIX and KNexPHENIX produce more substantial movements. ***C,*** In the KNexPHENIX-refined R220A metBJFIXL HEME domain crystal structure (PDB code 1Y28), a shift of R226 resolves a steric clash with T230 present in the original PDB structure. ***D,*** Steric hindrance is also avoided by rotation of the M156 side chain from T250 by KNexPHENIX refinement. ***E,*** PHENIX and REFMAC5 do not relieve a steric clash between L88 and T89 observed in the monoubiquitinated PCNA crystal structure (PDB code 3L0W) which is successfully resolved by KNexPHENIX.

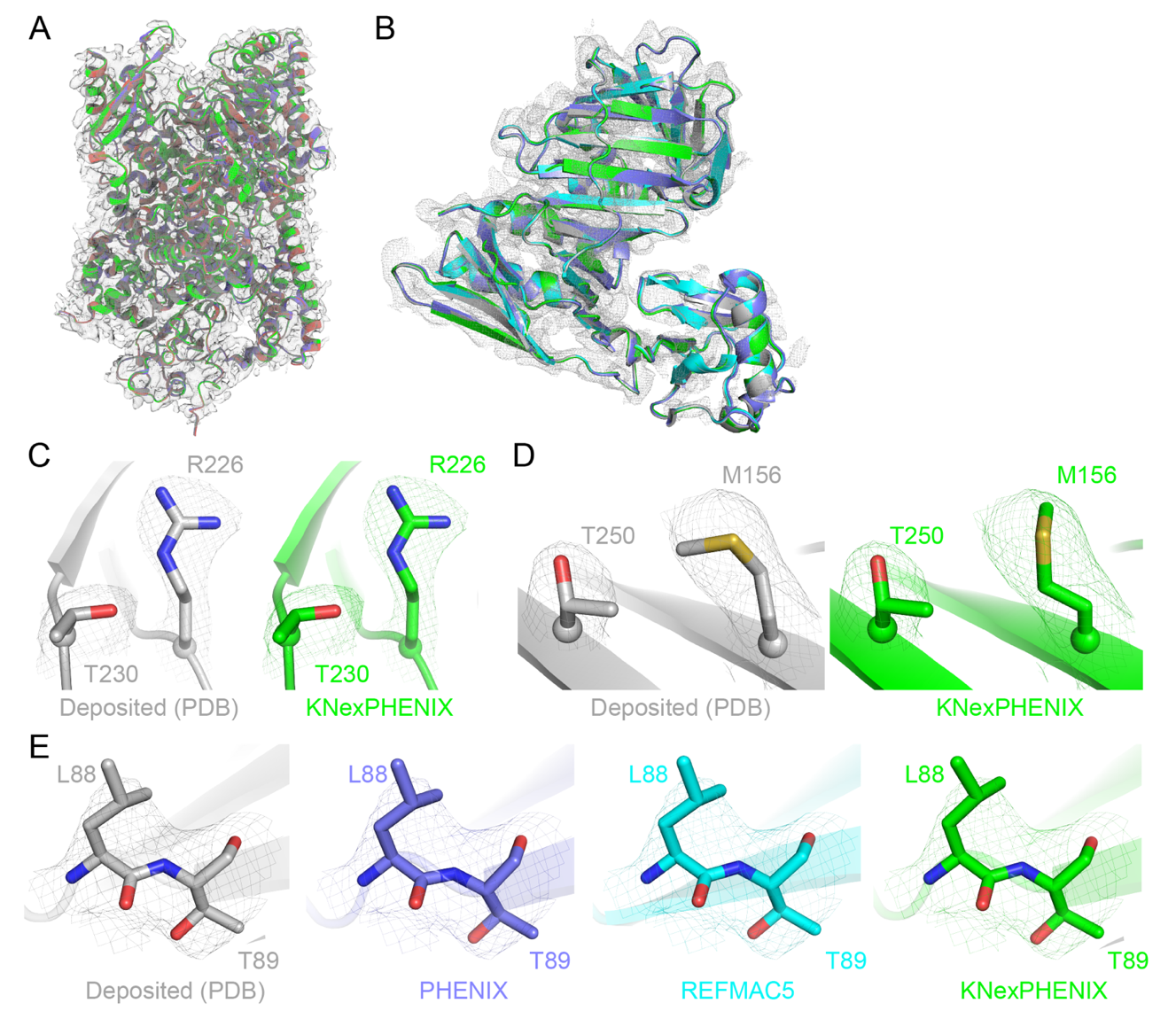

**Fig. S4. KNexPHENIX maintains a good model fit to map while improving model quality.** ***A,*** The PHENIX (blue), Servalcat (red), and KNexPHENIX (green) refined PI3Kalpha variant H1047R cryo-EM structure aligns well with the PDB model (gray) and fits well into the map. ***B,*** Similarly, monoubiquitinated PCNA refined by KNexPHENIX, PHENIX, REFMAC5 (cyan) also fits well into the map and is in overall good agreement with the deposited structure (PDB). KNexPHENIX refinement does not shift the residues significantly to affect map fit in the ***C-D,*** R220A metBJFIXL HEME domain crystal structure, or in ***E,*** the monoubiquitinated PCNA crystal structure. Related to **Fig. S3**.

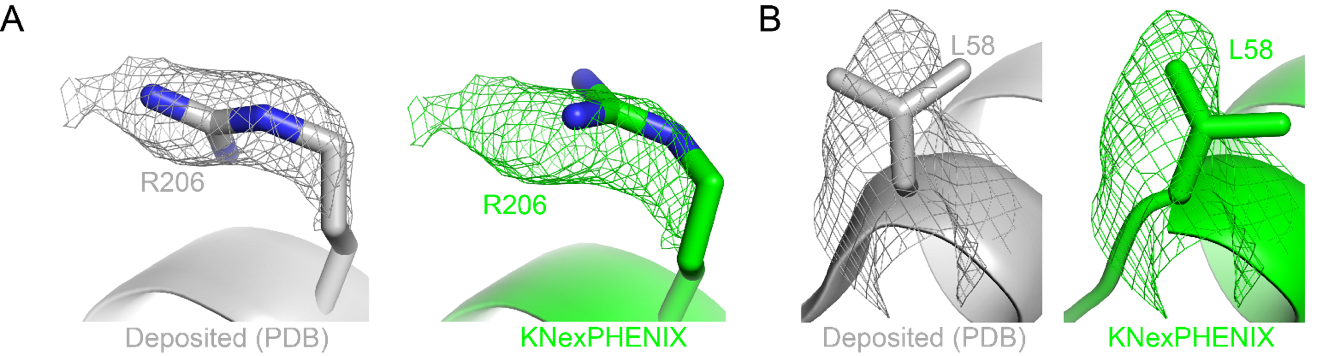

**Fig. S5. Overcorrection of rotamer outliers by KNexPHENIX results in poor model-to-map-fit. *A,*** KNexPHENIX (green) refinement of the deposited crystal structure (gray) of the metBJFIXL variant (PDB code 1Y28) results in correction of the rotamer outlier of R206 causing disagreement between model and map. ***B,*** Likewise, L58 in the deposited KDEL receptor variant structure (gray, PDB code 8APY) is well supported by the density, which is overcorrected by KNexPHENIX (green), shifting it outside the map.

**Supplementary Tables**

| **Table S1:** Structures used for KNexPHENIX benchmarking against other refinement approaches. | | | | | |
| --- | --- | --- | --- | --- | --- |
| PDB code | | Starting Model Use | Composition | Size (kDa) | Resolution (Å) |
| Cryo-EM | 9e0n | *De novo* only | Nucleoprotein complex | 2,280 | 3.24 |
|  | 5AN9 | Re-refine & *de novo* | Nucleoprotein complex | 1,419 | 3.30 |
|  | 8ETH | Re-refine & *de novo* | Nucleoprotein complex | 2,428 | 3.80 |
|  | 5H1S | Re-refine & *de novo* | Nucleoprotein complex | 1,445 | 3.50 |
|  | 6OF4 | Re-refine & *de novo* | Protein | 227 | 3.20 |
|  | 5A1A | Re-refine & *de novo* | Protein | 476 | 2.20 |
|  | 6JO5 | Re-refine & *de novo* | Protein | 765 | 2.90 |
|  | 8GUB | Re-refine & *de novo* | Protein | 212 | 2.73 |
|  | 8GUD | Re-refine & *de novo* | Protein | 128 | 2.62 |
|  | 6YEZ | Re-refine & *de novo* | Protein | 573 | 2.70 |
|  | 8ASW | Re-refine only | Nucleoprotein complex | 483 | 3.96 |
|  | 7UN3 | Re-refine only | Protein | 478 | 3.50 |
|  | 7W0L | Re-refine only | Protein | 223 | 3.57 |
|  | 7W0P | Re-refine only | Protein | 176 | 3.16 |
| X-ray crystallography | 1Y28 | Re-refine & *de novo* | Protein | 15 | 2.10 |
|  | 3L0W | Re-refine & *de novo* | Protein | 38 | 2.80 |
|  | 6KBI | Re-refine & *de novo* | Protein | 140 | 3.00 |
|  | 8F0V | Re-refine & *de novo* | Protein | 19 | 2.95 |
|  | 8HUK | Re-refine & *de novo* | Protein | 64 | 2.98 |
|  | 1YNS | Re-refine & *de novo* | Protein | 29 | 1.70 |
|  | 1FT2 | Re-refine & *de novo* | Protein | 83 | 3.40 |
|  | 1D8U | Re-refine & *de novo* | Protein | 38 | 2.35 |
|  | 6SZW | Re-refine & *de novo* | Protein-protein complex | 80 | 3.14 |
|  | 8APY | Re-refine & *de novo* | Protein-protein complex | 38 | 2.34 |
|  | 3UZ0 | Re-refine only | Protein-protein complex | 63 | 2.82 |
|  | 3JWR | Re-refine only | Protein-protein complex | 82 | 2.99 |
|  | 1ZOY | Re-refine only | Protein-protein complex | 127 | 2.40 |
|  | 4YJ5 | Re-refine only | Protein | 229 | 2.41 |
|  | 1R30 | Re-refine only | Protein | 85 | 3.40 |
|  | 1M52 | Re-refine only | Protein | 69 | 2.60 |

| **Table S2:** MolProbity score (MS) and CC_mask_ calculated from 13 cryo-EM structures deposited in the PDB and after their refinement using PHENIX, REFMAC Servalcat, and KNexPHENIX. | | | | | | | | |
| --- | --- | --- | --- | --- | --- | --- | --- | --- |
| PDB code | PDB (deposited)*^a^* | | PHENIX*^a^* | | REFMAC Servalcat*^a^* | | KNexPHENIX*^a^* | |
|  | MS | CC_mask_ | MS | CC_mask_ | MS | CC_mask_ | MS | CC_mask_ |
| 5AN9 | 2.02 | 0.81 | 2.38 | 0.80 | 3.17 | 0.84 | 1.79 | 0.78 |
| 8ETH | 1.89 | 0.61 | 2.20 | 0.64 | 3.04 | 0.70 | 1.52 | 0.61 |
| 6OF4 | 2.65 | 0.82 | 2.16 | 0.81 | 3.51 | 0.90 | 1.52 | 0.78 |
| 7UN3 | 2.00 | 0.63 | 2.03 | 0.68 | 3.61 | 0.80 | 1.50 | 0.62 |
| 8ASW | 2.35 | 0.78 | 2.77 | 0.78 | 3.81 | 0.88 | 1.73 | 0.74 |
| 7W0L | 2.15 | 0.71 | 2.31 | 0.78 | 3.61 | 0.88 | 1.77 | 0.74 |
| 5A1A | 1.97 | 0.76 | 2.28 | 0.79 | 1.82 | 0.80 | 1.32 | 0.76 |
| 8GUB | 2.35 | 0.65 | 2.45 | 0.78 | 2.82 | 0.82 | 1.24 | 0.76 |
| 6JO5 | 1.87 | 0.84 | 2.17 | 0.89 | 2.65 | 0.90 | 1.63 | 0.81 |
| 6YEZ | 2.35 | 0.87 | 2.17 | 0.91 | 2.72 | 0.93 | 1.93 | 0.89 |
| 5H1S | 3.24 | 0.77 | 2.80 | 0.82 | 3.34 | 0.89 | 2.33 | 0.77 |
| 8GUD | 2.20 | 0.56 | 2.14 | 0.54 | 3.72 | 0.64 | 1.50 | 0.53 |
| 7W0P | 2.05 | 0.59 | 2.24 | 0.70 | 3.14 | 0.83 | 1.64 | 0.67 |
| *Mean* | *2.24* | *0.72* | *2.32* | *0.76* | *3.15* | *0.83* | *1.65* | *0.73* |
| *^a^*Data used to generate plots shown in **Fig. 1A,B**. | | | | | | | | |

| **Table S3:** MolProbity score (MS) and CC_mask_ calculated from 10 cryo-EM structures deposited in the PDB and after their refinement using PHENIX, REFMAC Servalcat, CERES, and KNexPHENIX. | | | | | | | | | | |
| --- | --- | --- | --- | --- | --- | --- | --- | --- | --- | --- |
| PDB  code | PDB (deposited)*^a^* | | PHENIX*^a^* | | REFMAC Servalcat*^a^* | | CERES*^a^* | | KNexPHENIX*^a^* | |
|  | MS | CC_mask_ | MS | CC_mask_ | MS | CC_mask_ | MS | CC_mask_ | MS | CC_mask_ |
| 5AN9 | 2.02 | 0.81 | 2.38 | 0.80 | 3.17 | 0.84 | 2.25 | 0.81 | 1.79 | 0.78 |
| 8ETH | 1.89 | 0.61 | 2.20 | 0.64 | 3.04 | 0.70 | 1.86 | 0.60 | 1.52 | 0.61 |
| 6OF4 | 2.65 | 0.82 | 2.16 | 0.81 | 3.51 | 0.90 | 1.93 | 0.81 | 1.52 | 0.78 |
| 7UN3 | 2.00 | 0.63 | 2.03 | 0.68 | 3.61 | 0.80 | 1.97 | 0.69 | 1.50 | 0.62 |
| 8ASW | 2.35 | 0.78 | 2.77 | 0.78 | 3.81 | 0.88 | 2.21 | 0.78 | 1.73 | 0.74 |
| 7W0L | 2.15 | 0.71 | 2.31 | 0.78 | 3.61 | 0.88 | 2.01 | 0.77 | 1.77 | 0.74 |
| 5A1A | 1.97 | 0.76 | 2.28 | 0.79 | 1.82 | 0.80 | 1.95 | 0.79 | 1.32 | 0.76 |
| 8GUB | 2.35 | 0.65 | 2.45 | 0.78 | 2.82 | 0.82 | 2.00 | 0.79 | 1.24 | 0.76 |
| 5H1S | 3.24 | 0.77 | 2.80 | 0.82 | 3.72 | 0.64 | 2.41 | 0.83 | 2.33 | 0.77 |
| 8GUD | 2.20 | 0.56 | 2.14 | 0.54 | 3.14 | 0.83 | 2.26 | 0.56 | 1.50 | 0.53 |
| *Mean* | *2.28* | *0.71* | *2.35* | *0.74* | *3.23* | *0.81* | *2.09* | *0.74* | *1.62* | *0.71* |
| *^a^*Data used to generate plots shown in **Fig. 1C,D**. | | | | | | | | | | |

| **Table S4:** MolProbity score (MS) and CC_mask_ calculated from *de novo* refinement of 10 cryo-EM structures by PHENIX, REFMAC Servalcat, and KNexPHENIX using maps extracted from the PDB. | | | | | | | | |
| --- | --- | --- | --- | --- | --- | --- | --- | --- |
| PDB code*^a^* | | Starting Model | PHENIX*^b^* | | REFMAC Servalcat*^b^* | | KNexPHENIX*^b^* | |
|  |  |  | MS | CC_mask_ | MS | CC_mask_ | MS | CC_mask_ |
| 8gud | 8gua | | 2.22 | 0.50 | 3.39 | 0.61 | 1.17 | 0.45 |
| 8gub | 8dd4 | | 1.93 | 0.75 | 2.91 | 0.81 | 1.08 | 0.67 |
| 6of4 | 6of2 | | 2.09 | 0.79 | 3.45 | 0.89 | 1.29 | 0.74 |
| 5a1a | 6cvm | | 2.11 | 0.80 | 1.27 | 0.76 | 1.14 | 0.73 |
| 5h1s | 5x8t | | 2.66 | 0.83 | 3.26 | 0.89 | 1.94 | 0.78 |
| 6jo5 | 7bgi | | 1.39 | 0.86 | 1.96 | 0.88 | 0.78 | 0.79 |
| 8eth | 8ev3 | | 2.10 | 0.62 | 2.95 | 0.70 | 1.51 | 0.60 |
| 5an9 | 6qkl | | 2.64 | 0.83 | 3.17 | 0.85 | 1.76 | 0.80 |
| 6yez | 6zoo | | 1.71 | 0.92 | 2.24 | 0.92 | 0.90 | 0.86 |
| 9e0n | 5zeb | | 1.96 | 0.83 | 3.43 | 0.89 | 1.85 | 0.83 |
|  | *Mean* | | *2.08* | *0.77* | *2.80* | *0.82* | *1.34* | *0.73* |
| *^a^*Deposition corresponding to the map used for refinement.  *^b^*Data used to generate plots shown in **Fig. 2**. | | | | | | | | |

| **Table S5:** Comparison of MolProbity score (MS), R_work_, R_free_, ΔR (R_free_-R_work_), and/or CC_mask_ from *de novo* KNexPHENIX refinement of two cryo-EM and two crystal structures using a starting model from AlphaFold (KNexPHENIX-AF), Boltz2 (KNexPHENIX-Boltz2), RoseTTAFold3 (KNexPHENIX-RF3), and an initial model selected from the PDB. | | | | | | | | | | | | | | | | | | | | | |
| --- | --- | --- | --- | --- | --- | --- | --- | --- | --- | --- | --- | --- | --- | --- | --- | --- | --- | --- | --- | --- | --- |
| *Cryo-EM* | | | | | | | | | | | | | | | | | | | | | |
| PDB code*^a^* | KNexPHENIX*^b^* | | | | | KNexPHENIX-AF | | | | | | KNexPHENIX-Boltz2 | | | | KNexPHENIX-RF3 | | | | | |
|  | MS | | CC_mask_ | | | MS | | | CC_mask_ | | | MS | | CC_mask_ | | MS | | | CC_mask_ | | |
| 8GUD | 1.17 | | 0.45 | | | 0.77 | | | 0.44 | | | 0.87 | | 0.42 | | 0.96 | | | 0.42 | | |
| 5A1A*^c^* | 1.14 | | 0.73 | | | 1.04 | | | 0.74 | | | - | | - | | - | | | - | | |
| *X-ray Crystallography* | | | | | | | | | | | | | | | | | | | | | |
| PDB code*^a^* | KNexPHENIX*^b^* | | | | | | KNexPHENIX-AF | | | | | | KNexPHENIX-Boltz2 | | | | KNexPHENIX-RF3 | | | | |
|  | MS | R_work_ | | R_free_ | ΔR | | MS | R_work_ | | R_free_ | ΔR | | MS | R_work_ | R_free_ | ΔR | MS | R_work_ | | R_free_ | ΔR |
| 1Y28*^d^* | 0.91 | 28.4 | | 30.6 | 2.20 | | 1.29 | 29.7 | | 32.5 | 2.80 | | 1.39 | 31.9 | 32.8 | 0.90 | 1.17 | 33.5 | | 34.7 | 1.20 |
| 8F0V | 1.64 | 34.4 | | 39.0 | 4.60 | | 2.15 | 29.8 | | 40.4 | 10.6 | | 2.01 | 28.6 | 36.9 | 8.30 | 1.58 | 37.5 | | 41.9 | 4.40 |
| *^a^*Deposition corresponding to the map used for refinement.  *^b^*Original model selected from the PDB (see **Tables S4** and **S7**).  *^c^*Boltz2 and RF3 could not predict a starting model for 5A1A as the total number of residues is >3,500.  *^d^*The AF model corresponding to 1Y28 was truncated to remove residues without corresponding density prior to refinement. | | | | | | | | | | | | | | | | | | | | | |

| **Table S6:** MolProbity score (MS), R_work_, R_free_, and ΔR (R_free_-R_work_) calculated from 16 crystal structures deposited in the PDB and after their refinement using PHENIX, REFMAC5, and KNexPHENIX. | | | | | | | | | | | | | | | | |
| --- | --- | --- | --- | --- | --- | --- | --- | --- | --- | --- | --- | --- | --- | --- | --- | --- |
| PDB  code | PDB (deposited)*^a^* | | | | PHENIX*^a^* | | | | REFMAC5*^a^* | | | | KNexPHENIX*^a^* | | | |
|  | MS | R_work_ | R_free_ | ΔR | MS | R_work_ | R_free_ | ΔR | MS | R_work_ | R_free_ | ΔR | MS | R_work_ | R_free_ | ΔR |
| 1Y28 | 2.75 | 23.4 | 26.9 | 3.5 | 1.70 | 20.8 | 24.3 | 3.5 | 2.45 | 20.2 | 23.4 | 3.2 | 1.45 | 21.3 | 23.8 | 2.5 |
| 3UZ0 | 3.29 | 20.9 | 26.8 | 5.9 | 2.78 | 21.1 | 27.3 | 6.2 | 3.12 | 20.4 | 26.2 | 5.8 | 2.15 | 24.0 | 28.0 | 4.0 |
| 4YJ5 | 2.78 | 16.2 | 22.7 | 6.5 | 2.23 | 16.7 | 23.0 | 6.3 | 2.55 | 17.2 | 23.1 | 5.9 | 1.71 | 18.2 | 22.9 | 4.7 |
| 3L0W | 3.28 | 29.7 | 31.4 | 1.7 | 2.84 | 23.4 | 27.0 | 3.6 | 3.06 | 23.0 | 26.7 | 3.7 | 2.05 | 25.5 | 27.2 | 1.7 |
| 6SZW | 2.30 | 20.6 | 26.6 | 6.0 | 2.04 | 19.5 | 27.7 | 8.2 | 2.50 | 19.3 | 26.4 | 7.1 | 1.50 | 21.8 | 27.6 | 5.8 |
| 6KBI | 2.68 | 23.0 | 27.4 | 4.4 | 2.71 | 22.0 | 28.0 | 6.0 | 2.72 | 21.7 | 26.9 | 5.2 | 2.24 | 22.9 | 27.3 | 4.4 |
| 8APY | 2.28 | 27.2 | 32.8 | 5.6 | 2.63 | 25.9 | 33.6 | 7.7 | 2.02 | 26.5 | 32.3 | 5.8 | 1.50 | 28.4 | 33.1 | 4.7 |
| 8F0V | 2.82 | 25.7 | 29.2 | 3.5 | 3.42 | 25.1 | 31.9 | 6.8 | 3.16 | 24.9 | 30.0 | 5.1 | 2.10 | 28.1 | 31.4 | 3.3 |
| 8HUK | 2.49 | 23.0 | 26.7 | 3.7 | 2.92 | 20.8 | 28.7 | 7.9 | 2.32 | 21.1 | 26.9 | 5.8 | 1.72 | 23.0 | 26.4 | 3.4 |
| 1R30 | 3.80 | 20.6 | 24.6 | 4.0 | 2.53 | 18.4 | 24.5 | 6.1 | 3.47 | 18.0 | 23.9 | 5.9 | 2.13 | 21.1 | 25.1 | 4.0 |
| 1YNS | 2.82 | 21.0 | 21.5 | 0.5 | 2.00 | 20.2 | 23.8 | 3.6 | 2.50 | 18.1 | 21.4 | 3.3 | 2.07 | 19.8 | 24.6 | 4.8 |
| 3JWR | 3.19 | 20.9 | 27.7 | 6.8 | 2.60 | 19.1 | 27.5 | 8.4 | 2.87 | 20.7 | 27.8 | 7.1 | 1.99 | 22.0 | 26.7 | 4.7 |
| 1FT2 | 2.68 | 21.8 | 25.9 | 4.1 | 2.32 | 19.4 | 26.2 | 6.8 | 2.68 | 20.4 | 26.1 | 5.7 | 1.70 | 22.1 | 25.5 | 3.4 |
| 1D8U | 2.63 | 21.0 | 21.0 | 0.0 | 2.24 | 20.1 | 25.8 | 5.7 | 2.05 | 19.9 | 24.3 | 4.4 | 1.93 | 21.0 | 24.8 | 3.8 |
| 1M52 | 2.62 | 20.3 | 24.7 | 4.4 | 2.10 | 18.0 | 24.1 | 6.1 | 2.28 | 18.7 | 24.1 | 5.4 | 1.99 | 18.4 | 24.1 | 5.7 |
| 1ZOY | 3.24 | 20.5 | 25.2 | 4.7 | 2.88 | 19.5 | 25.4 | 5.9 | 3.00 | 19.2 | 24.7 | 5.5 | 2.31 | 21.5 | 25.4 | 3.9 |
| *Mean* | *2.85* | *22.3* | *26.2* | *4.0* | *2.50* | *20.6* | *26.8* | *6.2* | *2.67* | *20.6* | *25.9* | *5.3* | *1.91* | *22.4* | *26.5* | *4.1* |
| *^a^*Data used to generate plots shown in **Fig. 3**. | | | | | | | | | | | | | | | | |

| **Table S7:** MolProbity score (MS), R_work_, R_free_, and ΔR (R_free_-R_work_) calculated from *de novo* refinement of 10 crystal structures by PHENIX, REFMAC5, and KNexPHENIX using maps extracted from the PDB. | | | | | | | | | | | | | |
| --- | --- | --- | --- | --- | --- | --- | --- | --- | --- | --- | --- | --- | --- |
| PDB code*^a^* | Starting Model*^b^* | PHENIX*^c^* | | | | REFMAC5*^c^* | | | | KNexPHENIX*^c^* | | | |
|  |  | MS | R_work_ | R_free_ | ΔR | MS | R_work_ | R_free_ | ΔR | MS | R_work_ | R_free_ | ΔR |
| 1YNS | 1ZS9 | 1.09 | 26.4 | 28.5 | 2.1 | 1.66 | 26.5 | 28.0 | 1.5 | 0.94 | 26.9 | 28.6 | 1.7 |
| 3L0W | 1PLQ | 2.79 | 25.8 | 29.6 | 3.8 | 2.65 | 25.0 | 27.2 | 2.2 | 1.45 | 29.8 | 30.5 | 0.7 |
| 1Y28 | 1DRM | 1.95 | 27.1 | 30.0 | 2.9 | 2.08 | 27.1 | 29.3 | 2.2 | 0.91 | 28.4 | 30.6 | 2.2 |
| 1FT2 | 1FT1 | 2.39 | 19.7 | 26.1 | 6.4 | 2.43 | 21.0 | 25.9 | 4.9 | 1.06 | 25.1 | 27.8 | 2.7 |
| 1D8U | 2GNW | 2.23 | 25.5 | 30.3 | 4.8 | 2.67 | 25.8 | 29.1 | 3.3 | 1.46 | 27.0 | 30.5 | 3.5 |
| 6SZW | 1P32 | 2.15 | 25.8 | 32.2 | 6.4 | 1.94 | 26.6 | 31.0 | 4.4 | 0.97 | 30.2 | 32.0 | 1.8 |
| 6KBI | 1M6B | 2.82 | 23.2 | 28.5 | 5.3 | 3.04 | 23.1 | 27.3 | 4.2 | 1.51 | 26.2 | 29.1 | 2.9 |
| 8APY | 6I6J | 2.54 | 28.4 | 34.7 | 6.3 | 1.85 | 29.4 | 33.8 | 4.4 | 1.58 | 31.3 | 35.0 | 3.7 |
| 8F0V | 7BKX | 3.08 | 28.5 | 36.4 | 8.1 | 3.32 | 27.5 | 33.6 | 6.1 | 1.64 | 34.4 | 39.0 | 4.6 |
| 8HUK | 3SP6 | 2.80 | 24.7 | 32.1 | 7.4 | 2.18 | 25.1 | 29.6 | 4.5 | 1.54 | 28.9 | 32.4 | 3.5 |
|  | *Mean* | *2.38* | *25.5* | *30.9* | *5.3* | *2.38* | *25.7* | *29.5* | *3.8* | *1.31* | *28.8* | *31.6* | *2.7* |
| *^a^*Deposition corresponding to the map used for refinement.  *^b^*Model used for molecular replacement (MR).  *^c^*Data used to generate plots shown in **Fig. 4**. | | | | | | | | | | | | | |

| **Table S8**: Clashscores calculated from deposited crystal structures and de novo refinement using PHENIX and KNexPHENIX. | | | | |
| --- | --- | --- | --- | --- |
| PDB model | Deposited | | De novo | |
|  | PHENIX | KNexPHENIX | PHENIX | KNexPHENIX |
| 1Y28 | 6.20 | 3.60 | 5.40 | 1.60 |
| 3UZ0 | 13.1 | 8.60 | - | - |
| 4YJ5 | 8.40 | 5.90 | - | - |
| 3L0W | 13.6 | 9.20 | 10.6 | 4.40 |
| 6SZW | 9.50 | 3.50 | 13.5 | 1.80 |
| 6KBI | 13.7 | 8.90 | 13.0 | 3.80 |
| 8APY | 14.1 | 7.90 | 11.1 | 4.20 |
| 8F0V | 55.0 | 10.2 | 29.0 | 8.00 |
| 8HUK | 15.6 | 8.80 | 16.8 | 10.4 |
| 1R30 | 28.6 | 15.4 | - | - |
| 1YNS | 23.3 | 27.3 | 1.80 | 1.80 |
| 3JWR | 11.6 | 9.30 | - | - |
| 1FT2 | 18.0 | 7.90 | 19.9 | 2.70 |
| 1D8U | 9.70 | 7.30 | 7.90 | 3.50 |
| 1M52 | 8.40 | 6.30 | - | - |
| 1ZOY | 20.8 | 12.1 | - | - |
| *Mean* | *16.8* | *9.50* | *12.9* | *4.20* |

| **Table S9**: Clashscores calculated from deposited cryo-EM structures and de novo refinement using PHENIX and KNexPHENIX. | | | | |
| --- | --- | --- | --- | --- |
| PDB model | Deposited | | De novo | |
|  | PHENIX | KNexPHENIX | PHENIX | KNexPHENIX |
| 5AN9 | 11.7 | 6.10 | 14.3 | 6.70 |
| 8ETH | 11.4 | 5.40 | 14.1 | 6.20 |
| 6OF4 | 7.60 | 2.50 | 8.30 | 1.40 |
| 7UN3 | 13.6 | 6.20 | - | - |
| 8ASW | 18.2 | 6.10 | - | - |
| 7W0L | 12.0 | 6.80 | - | - |
| 5A1A | 12.9 | 5.90 | 8.60 | 3.10 |
| 8GUB | 33.4 | 4.70 | 10.3 | 2.30 |
| 6JO5 | 15.3 | 9.20 | 4.50 | 0.90 |
| 6YEZ | 14.9 | 12.2 | 5.80 | 1.60 |
| 5H1S | 15.0 | 11.3 | 13.9 | 7.60 |
| 8GUD | 16.1 | 5.90 | 14.3 | 2.00 |
| 7W0P | 13.0 | 5.40 | - | - |
| 9E0N | - | - | 17.2 | 8.50 |
| *Mean* | *15.0* | *6.70* | *11.1* | *4.00* |

**Supplementary Methods**

**Default PHENIX refinement for cryo-EM and X-ray crystal structures**

For performance comparisons in refinement of cryo-EM structures with KNexPHENIX, five cycles of “default” PHENIX refinement (*phenix.real_space_refine*) were performed using local grid search, global minimization, occupancy, N/Q/H flips, and B-factor refinement, with target root mean square deviation (RMSD) of 0.01Å and 1.0˚ for bonds and angles, respectively. Restraints such as secondary structure and Ramachandran were applied but not reference model restraints. For *pdb_interpretation*, Ramachandran restraints for only peptide bonds were added, but peptide planarity constraints were not employed. The dihedral function type was set to be determined by the sign of periodicity. Similarly, unless stated otherwise, for X-ray crystallographic structure refinement comparisons, five cycles of PHENIX refinement (*phenix.refine*) were performed using real-space, reciprocal-space, occupancy, and B-factor refinement. For *pdb_interpretation*, the dihedral function type was set to be determined by the sign of periodicity. For all the steps, parameters not described above were set to their default values. All the parameters in the two refinement procedures described here employed default values for parameters not described here.

**Model refinement in REFMAC**

Cryo-EM structures were refined using twenty cycles of REFMAC Servalcat (version 1.6.0) masked refinement with the weight and symmetry chosen automatically, and addition of hydrogen atoms. No RNA/DNA restraints were added, sharpening was not performed, and jellybody refinement was turned off. Crystal structures were refined in REFMAC5 (version 5.8.0419) through ten cycles of maximum likelihood restrained refinement using default parameters, e.g., isotropic B-factors, automatically optimized and experimental sigma stereochemistry/X-ray weights, and hydrogen atom addition. MolProbity scores^1^, CC_mask_^2^, and R_work_/R_free_^3^ values for the REFMAC-refined cryo-EM and crystal structures were calculated using the PHENIX validation tool. The MolProbity scores, R_work_, and R_free_ values for the crystal structures deposited in PDB were also calculated similarly. Any options not explicitly mentioned were kept at their default values throughout.

**Supplementary Results**

**Motivation for stage and parameter selection in the KNexPHENIX refinement pipeline**

Selection of the stages and parameters in KNexPHENIX refinement was based on a defined rationale, although certain parameters were optimized empirically through trial and error. The guiding principle of the refinement protocol is to maximize the model quality while maintaining an appropriate map-to-model fit. The first step of the refinement pipeline involves the addition of H-atoms, which was suggested to improve the model geometry for crystal structures by correcting Asn/Gln/His flips and reducing clashes^1,4,5^. For the de novo refinement of crystal and cryo-EM structures, the next step involved refinement with simulated annealing, among other strategies, which improves agreement with the experimental map. Additionally, restraints on secondary structure, stereochemistry, reference model, etc., were applied to maintain chemically reasonable geometry. Although the PHENIX refinement of deposited cryo-EM models was observed to enhance the final KNexPHENIX refined model, we omitted this step for the crystal structures, as it did not appreciably alter the quality of the structure.

The subsequent step in the pipeline is geometry minimization, which is known to improve stereochemical parameters^6^. The number of minimization cycles was altered depending on (i) the confidence in the starting model quality, and (ii) eventual map-to-model fit. Deposited structures were subjected to a lower number of cycles (1-2) due to their extensive prior refinement and manual model corrections, thereby requiring modest improvements to improve their suitability for PDB deposition. In contrast, de novo models were passed through multiple rounds of minimization (2-5 cycles) to improve the model quality. However, since the minimization process is independent of the model and therefore carries a risk of overcorrection, it was followed by an additional round of PHENIX refinement focusing on fitting the model to the map. This refinement was accompanied by several restraints, particularly the harmonic restraints on the starting coordinates, in addition to the ones present in the first PHENIX refinement, to prevent excessive deviations from the improved geometry.

All in all, the sequence of steps in KNexPHENIX is designed to enhance model quality while maintaining consistency with the experimental data, thereby producing a refined structure suitable for immediate deposition.

**Typical duration and efficiency of KNexPHENIX refinement**

The total runtime for the KNexPHENIX workflow depends on the molecular weight of the structure, with larger molecules requiring a longer time. To provide an estimate of the time required, we report the wall-clock time for the KNexPHENIX refinements (de novo and deposited) for two representative structures spanning the lowest and highest molecular weights obtained from cryo-EM and X-ray crystallography, respectively. The de novo refinement of crystal structures with PDB codes 1Y28 (15 kDa) and 6KBI (140 kDa) required 32.6 and 245 min, respectively. Similarly, the time required for re-refining deposited structures 1Y28 and 4YJ5 (229 kDa) was 11.3 and 72 min, respectively. For the cryo-EM structures 8GUD (15 kDa) and 8ETH (2,428 kDa), the de novo refinement takes 88 and 764 min, respectively, and 11.3 and 72 min, respectively, for re-refining the deposited structure.

The ReadySet step does not report the wall-clock time, and therefore, it was excluded from these estimates. Additionally, for the de novo refinement of the cryo-EM structures, the initial docking of the model into the map was performed manually and therefore could not be timed. Similarly, the removal of the H-atoms was also conducted manually via the command line interface and was not included in the timing analysis.

With respect to human intervention, steps such as PHASER, ReadySet, PHENIX refinement, and geometry minimization require manual input of the model or both the map and the model. Also, as noted above, docking of de novo models into cryo-EM maps and removal of H-atoms are manual processes. In contrast, default PHENIX typically requires the user to provide the model and map only once.

Regarding the efficiency of the KNexPHENIX protocol compared to the default approach, the latter often requires a substantial amount of time to manually correct the geometric outliers of the model. The time required for that is highly speculative, as it depends on user experience as well as the structural complexity. In contrast, KNexPHENIX requires minimal manual intervention in correcting geometric outliers while ensuring map-to-model fit and therefore has the potential to save time ranging from several days to several months. We have included these details in a separate supplementary file and added a sentence in the manuscript referring to the information.

**Analyses of the effect of variation of stages and parameters in KNexPHENIX**

The different steps and specific criteria in KNexPHENIX were varied in the de novo refinement of two crystal and cryo-EM structures (modified KNexPHENIX is referred to as KNexPHENIX-M) to highlight their importance. The structures were chosen to represent the range in size (low and high molecular weight), resolution (poor and better), and type of molecule (single molecule *vs.* complex). The MolProbity score (MS) and map-to-model fit parameters for specific parameter variations are listed below.

1. ***No reference model restraints (RMR) in the final PHENIX refinement for cryo-EM structures.*** These analyses used 6OF4 (3.20 Å, 227 kDa, protein), 8GUD (2.60 Å, 128 kDa, protein), and 5AN9 (3.30 Å, 1,419 kDa, nucleoprotein complex). Additionally, the reference coordinate restraints (RCR) were also removed in *pdb_interpretation.* Absence of the restraints affected the MolProbity score as the improvements in the model from the geometry minimization in the previous step were partially nullified.

| PDB model*^a^* | Starting model*^b^* | KNexPHENIX | |  | | KNexPHENIX-M | | |
| --- | --- | --- | --- | --- | --- | --- | --- | --- |
|  |  | MS | CC_mask_ |  | Modification | | MS | CC_mask_ |
| 6OF4 | 6of2 | 1.29 | 0.74 |  | No RMR | | 1.42 | 0.76 |
|  |  |  |  |  | No RMR+RCR | | 1.23 | 0.45 |
| 8GUD | 8gua | 1.17 | 0.45 |  | No RMR | | 2.17 | 0.77 |
|  |  |  |  |  | No RMR+RCR | | 2.13 | 0.50 |
| 5AN9 | 6qkl | 1.76 | 0.80 |  | No RMR | | 2.01 | 0.80 |
|  |  |  |  |  | No RMR+RCR | | 2.77 | 0.82 |

*^a^*Deposition corresponding to the map used for refinement.

*^b^*Model used for molecular replacement (MR).

2. ***Absence of simulated annealing (SA) in the final PHENIX refinement of crystal structures.*** These analyses used 1Y28 (2.10 Å, 15 kDa, protein), 8F0V (2.95 Å, 19 kDa, protein), and 6KBI (3.00 Å, 140 kDa, protein). As an alternative parameter variation, geometry restraints (GR) were also removed in the final step. In 1Y28, no SA reduced the map-to-model fit and thereby increased the R_work_ and R_free_. On the other hand, removal of GR reduces the MolProbity score by affecting model quality. Interestingly, absence of SA or GR only affects the MolProbity score in 8F0V and 6KBI, reasons for which are not clearly understood.

| PDB model*^a^* | Starting model*^b^* | KNexPHENIX | | | |  |  | KNexPHENIX-M | | | |
| --- | --- | --- | --- | --- | --- | --- | --- | --- | --- | --- | --- |
|  |  | MS | R_work_ | R_free_ | ΔR*^c^* |  | Modification | MS | R_work_ | R_free_ | ΔR*^c^* |
|  |  |  |  |  |  |  | No SA | 0.91 | 29.3 | 31.5 | 2.20 |
| 1Y28 | 1DRM | 0.91 | 28.4 | 30.6 | 2.20 |  |  |  | | | |
|  |  |  |  |  |  |  | No GR | 1.06 | 28.3 | 30.6 | 2.30 |
|  |  |  |  |  |  |  | No SA | 1.82 | 34.5 | 38.3 | 3.80 |
| 8F0V | 7BKX | 1.64 | 34.4 | 39 | 4.60 |  |  |  | | | |
|  |  |  |  |  |  |  | No GR | 1.97 | 31.1 | 36.7 | 5.60 |
|  |  |  |  |  |  |  | No SA | 1.84 | 26.2 | 29.3 | 3.10 |
| 6KBI | 1M6B | 1.51 | 26.2 | 29.1 | 2.90 |  |  |  | | | |
|  |  |  |  |  |  |  | No GR | 1.68 | 24.8 | 28.6 | 3.80 |

*^a^*Deposition corresponding to the map used for refinement. *^b^*Model used for molecular replacement (MR). *^c^*ΔR is R_free_-R_work_

3. ***Omission of final PHENIX refinement for the cryo-EM structures***. Absence of the refinement affects both model quality and agreement with the map highlighting the importance of the refinement strategies and geometry restraints in this step.

| PDB model*^a^* | Starting model*^b^* | KNexPHENIX | |  | KNexPHENIX-M | |
| --- | --- | --- | --- | --- | --- | --- |
|  |  | MS | CC_mask_ |  | MS | CC_mask_ |
| 6OF4 | 6OF2 | 1.29 | 0.74 |  | 1.43 | 0.72 |
| 8GUD | 8GUA | 1.17 | 0.45 |  | 1.28 | 0.43 |
| 5AN9 | 6QKL | 1.76 | 0.80 |  | 1.83 | 0.77 |

*^a^*Deposition corresponding to the map used for refinement.

*^b^*Model used for molecular replacement (MR).

4. ***Removal of geometry minimization from the KNexPHENIX pipeline for crystal structures.*** As expected, failure to minimize the structure led to a considerable increase in the MolProbity scores.

| PDB model*^a^* | Starting model*^b^* | KNexPHENIX | | | |  |  | KNexPHENIX-M | | | |
| --- | --- | --- | --- | --- | --- | --- | --- | --- | --- | --- | --- |
|  |  | MS | R_work_ | R_free_ | ΔR*^c^* |  |  | MS | R_work_ | R_free_ | ΔR*^c^* |
| 1Y28 | 1DRM | 0.91 | 28.4 | 30.6 | 2.20 |  |  | 1.58 | 26.9 | 29.6 | 2.70 |
| 8F0V | 7BKX | 1.64 | 34.4 | 39.0 | 4.60 |  |  | 2.74 | 31.7 | 34.8 | 3.10 |
| 6KBI | 1M6B | 1.51 | 26.2 | 29.1 | 2.90 |  |  | 2.75 | 24.7 | 27.7 | 3.00 |

*^a^*Deposition corresponding to the map used for refinement. *^b^*Model used for molecular replacement (MR). *^c^*ΔR is R_free_-R_work_

In conclusion, the alterations in parameters and stages successfully demonstrate their importance in the KNexPHENIX pipeline to obtain a model suitable for deposition.

6.	phenix.geometry_minimization: regularize model geometry. <https://www.phenix-online.org/version_docs/dev-2486/reference/geometry_minimization.html> (accessed 01/04).

**Case Study 1: PI3Kalpha H1047R cryo-EM structure (PDB code 8GUB)**

Changes in validation parameters during each stage of *de novo* refinement of PI3Kalpha H1047R variant cryo-EM structure using **KNexPHENIX Workflow 2**.

**PHENIX refinement**

| ***Refinement and model*** |  |
| --- | --- |
| Model resolution (FSC 0.143, unmasked), Å | 2.7 |
| CC_mask_ | 0.67 |
| RMS deviations |  |
| Bond lengths, Å | 0.002 |
| Bond angles, ° | 0.545 |
| ***Validation*** |  |
| MolProbity score | 1.54 |
| Clashscore | 6.73 |
| Rotamer outliers, % | 0.00 |
| Ramachandran plot (protein) |  |
| Favored, % | 97.0 |
| Allowed, % | 2.8 |
| Disallowed, % | 0.2 |

**Geometry minimization**

| ***Refinement and model*** |  |
| --- | --- |
| RMS deviations |  |
| Bond lengths, Å | 0.001 |
| Bond angles, ° | 0.322 |
| ***Validation*** |  |
| Clashscore | 3.03 |
| Rotamer outliers, % | 0.26 |
| Ramachandran plot (protein) |  |
| Favored, % | 97.6 |
| Allowed, % | 2.3 |
| Disallowed, % | 0.1 |

**PHENIX refinement**

| ***Refinement and model*** |  |
| --- | --- |
| Model resolution (FSC 0.143, unmasked), Å | 2.7 |
| CC_mask_ | 0.67 |
| RMS deviations |  |
| Bond lengths, Å | 0.002 |
| Bond angles, ° | 0.426 |
| ***Validation*** |  |
| MolProbity score | 1.08 |
| Clashscore | 2.26 |
| Rotamer outliers, % | 0.26 |
| Ramachandran plot (protein) |  |
| Favored, % | 97.6 |
| Allowed, % | 2.3 |
| Disallowed, % | 0.8 |

**Case study 2**: **Monoubiquitinated PCNA X-ray crystal structure (PDB code 3L0W)**

Changes in validation parameters during each stage of *de novo* refinement of monoubiquitinated PCNA crystal structure using **KNexPHENIX Workflow 4**.

**PHENIX refinement**

| ***Refinement and model*** |  |
| --- | --- |
| R_work_ (%) | 27.45 |
| R_free_ (%) | 28.17 |
| R_free_-R_work_ (%) | 0.72 |
| RMS deviations |  |
| Bond lengths, Å | 0.013 |
| Bond angles, ° | 1.836 |
| ***Validation*** |  |
| MolProbity score | 2.39 |
| Clashscore | 19.16 |
| Rotamer outliers, % | 1.72 |
| Ramachandran plot (protein) |  |
| Favored, % | 93.4 |
| Allowed, % | 4.7 |
| Disallowed, % | 1.9 |

**Geometry minimization**

| ***Refinement and model*** |  |
| --- | --- |
| RMS deviations |  |
| Bond lengths, Å | 0.001 |
| Bond angles, ° | 0.384 |
| ***Validation*** |  |
| Clashscore | 2.70 |
| Rotamer outliers, % | 0.86 |
| Ramachandran plot (protein) |  |
| Favored, % | 96.9 |
| Allowed, % | 3.1 |
| Disallowed, % | 0.0 |

**PHENIX refinement**

| ***Refinement and model*** |  |
| --- | --- |
| R_work_ (%) | 29.76 |
| R_free_ (%) | 30.53 |
| R_free_-R_work_ (%) | 0.77 |
| RMS deviations |  |
| Bond lengths, Å | 0.010 |
| Bond angles, ° | 1.168 |
| ***Validation*** |  |
| MolProbity score | 1.45 |
| Clashscore | 4.42 |
| Rotamer outliers, % | 0.86 |
| Ramachandran plot (protein) |  |
| Favored, % | 96.5 |
| Allowed, % | 3.5 |
| Disallowed, % | 0.0 |

**KNexPHEX “How-to” guide (Workflows 1-4 with PHENIX screenshots)**

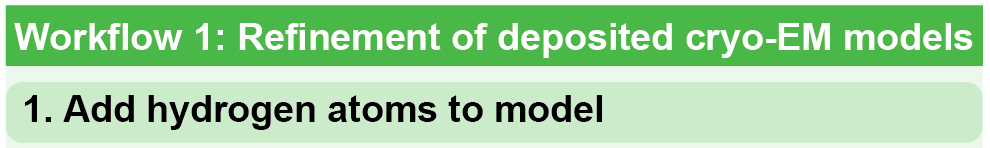

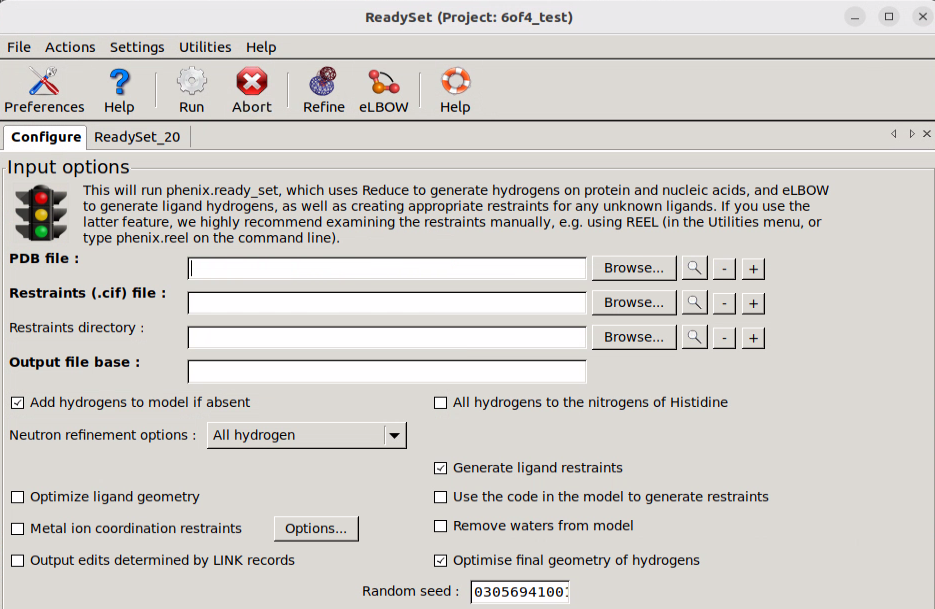

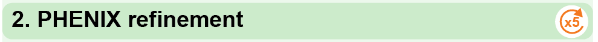

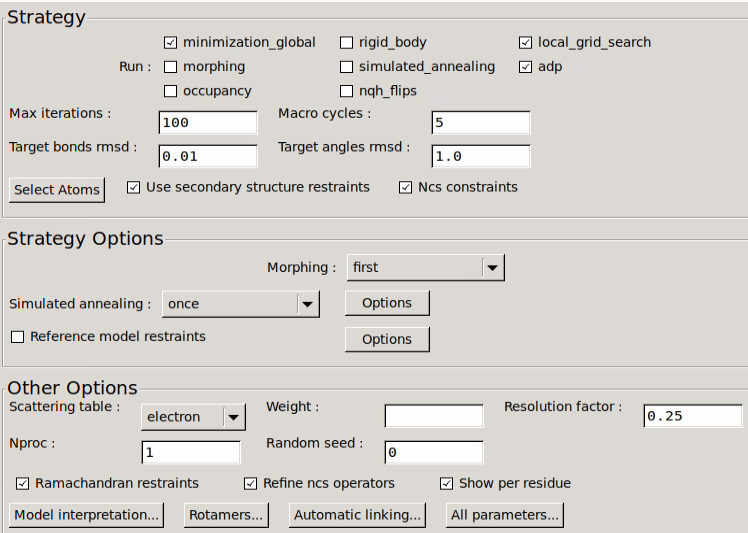

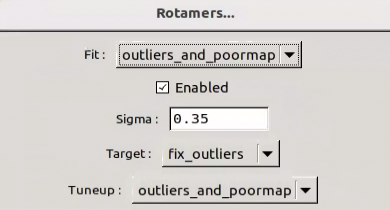

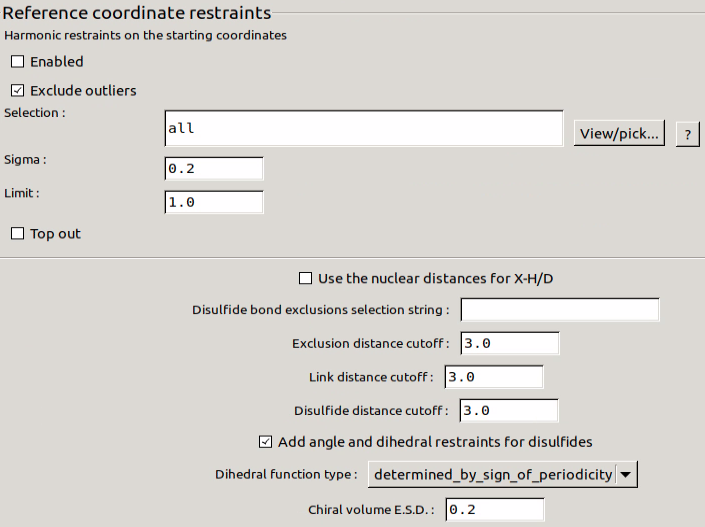

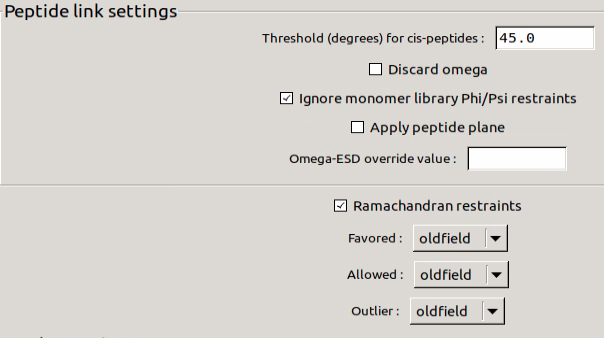

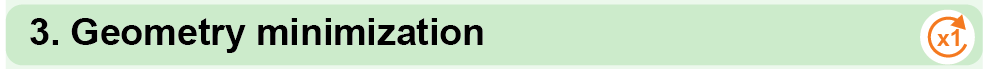

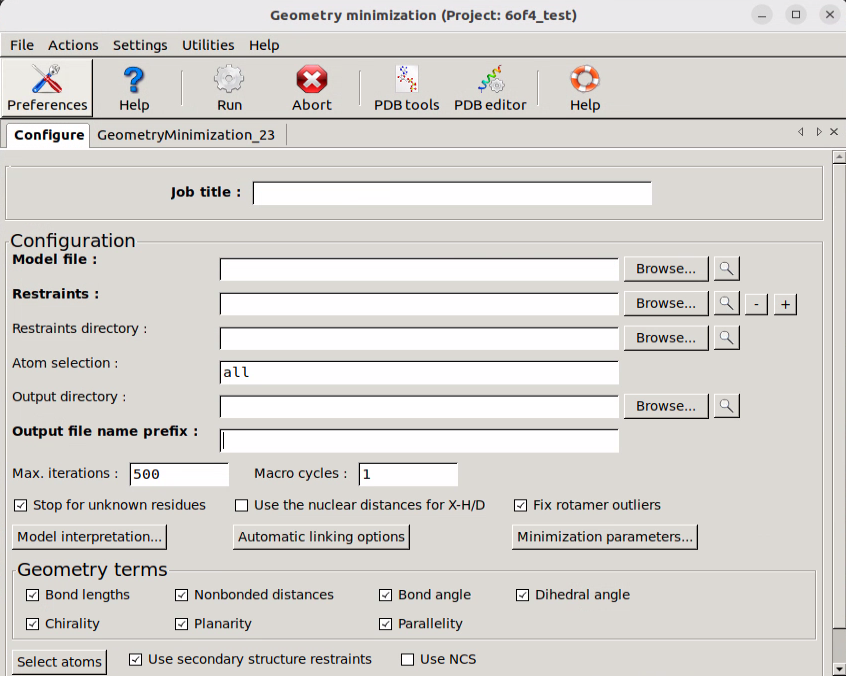

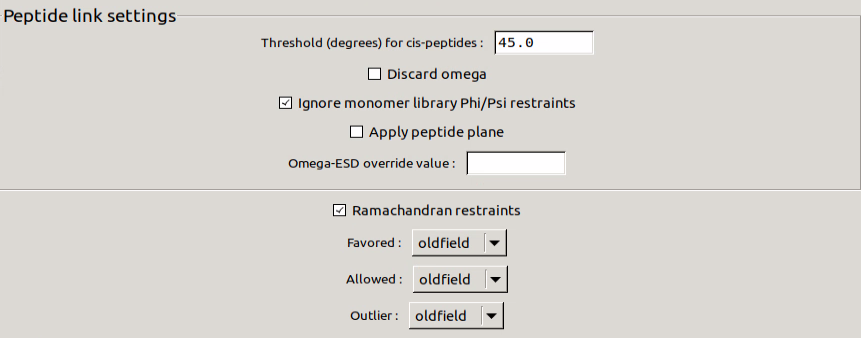

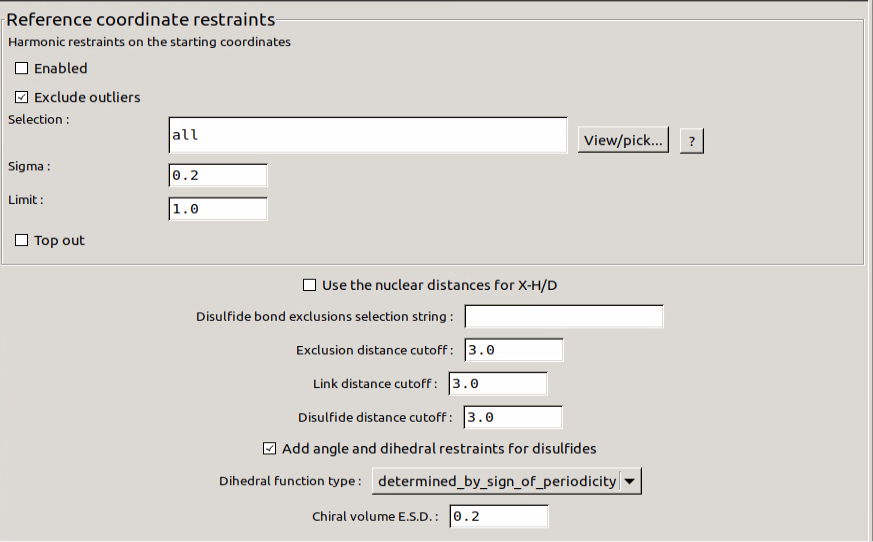

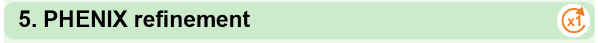

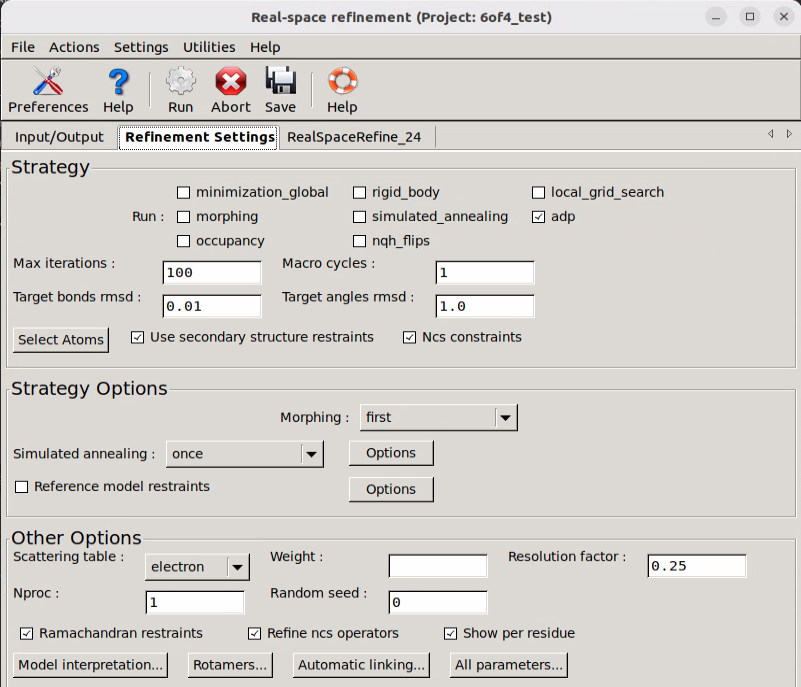

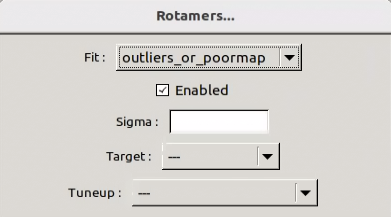

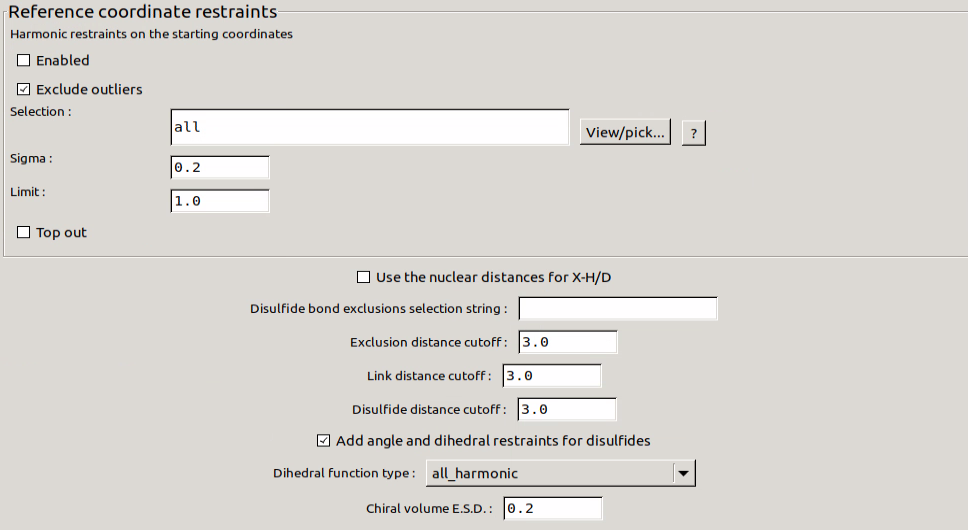

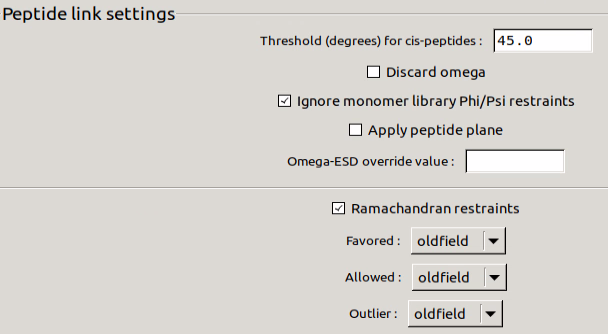

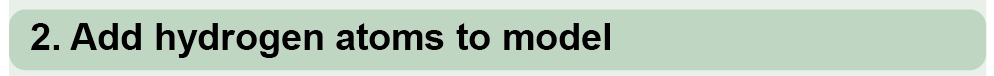

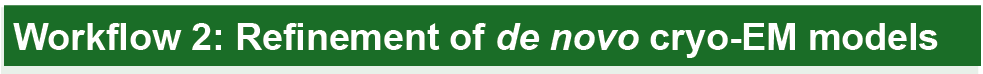

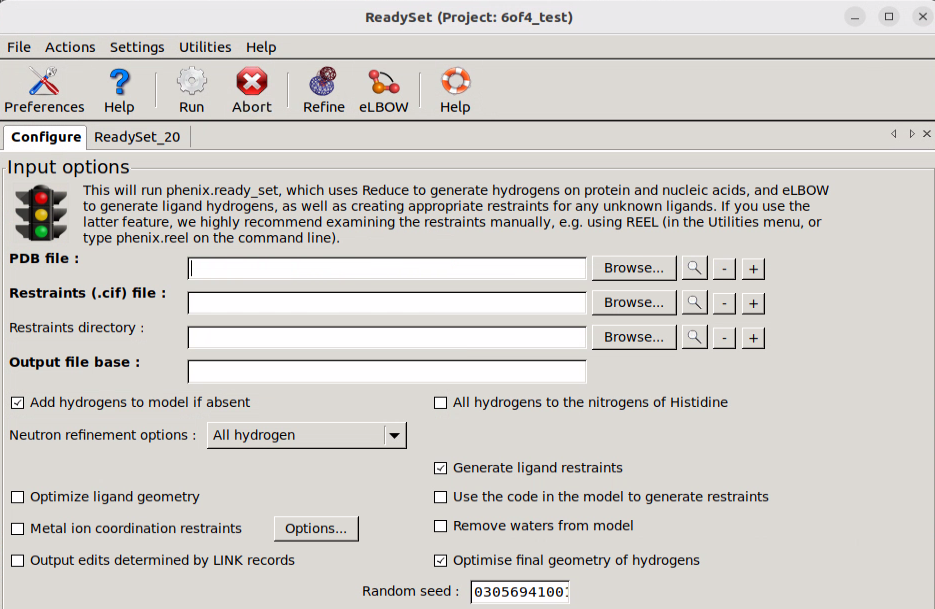

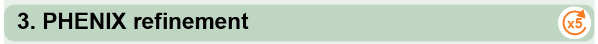

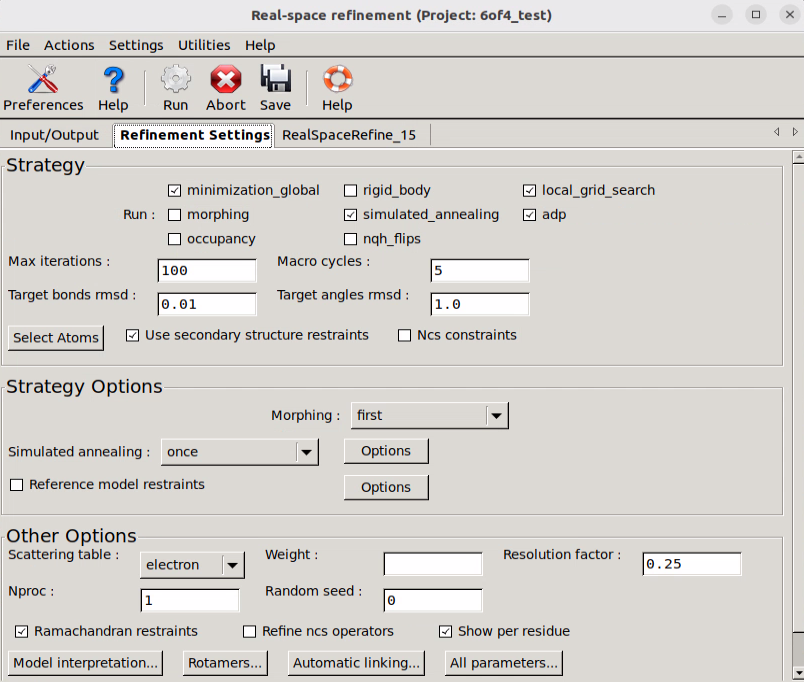

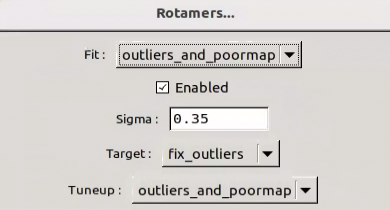

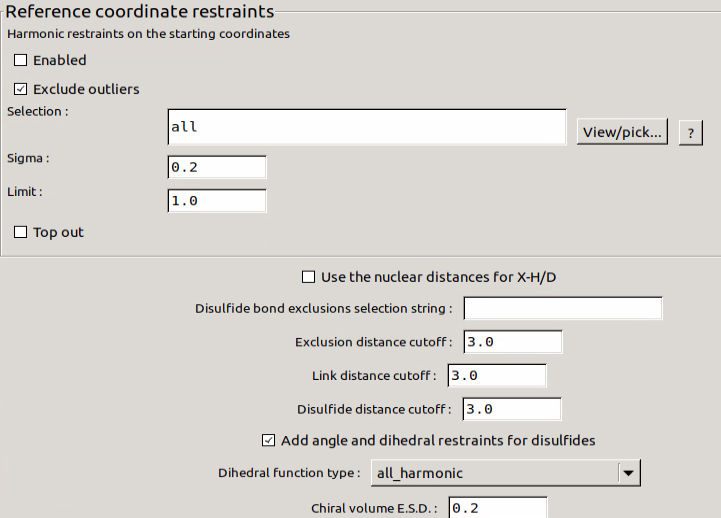

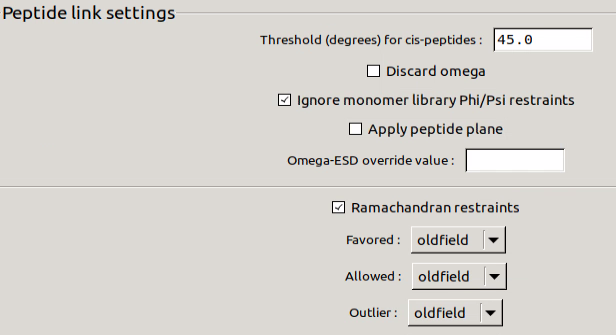

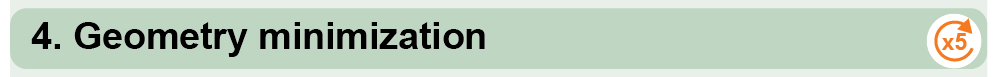

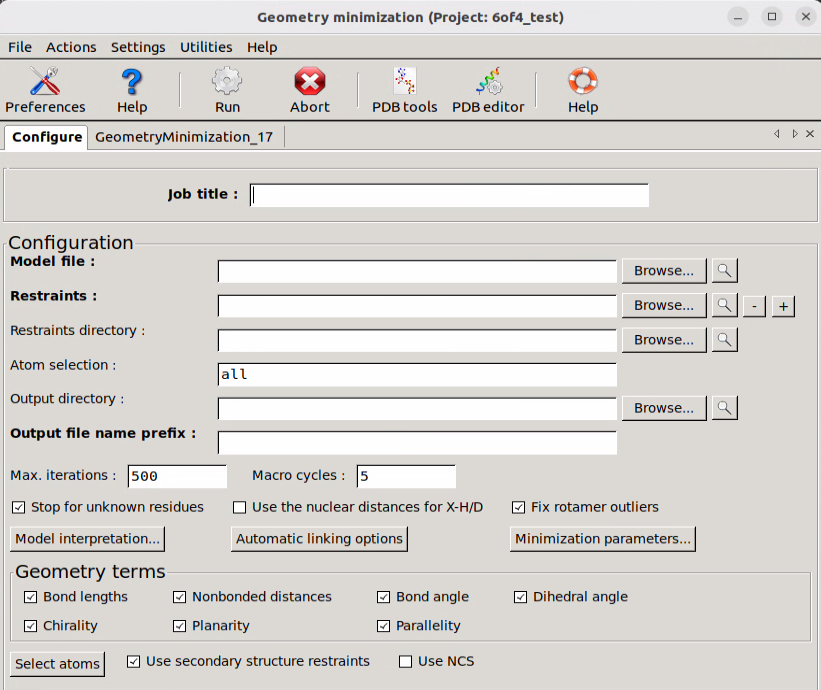

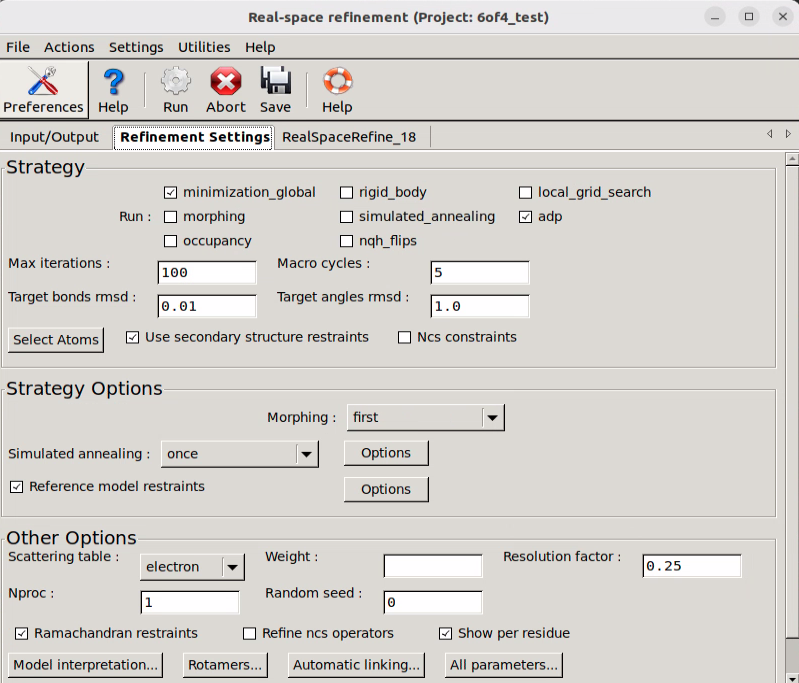

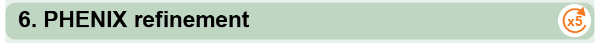

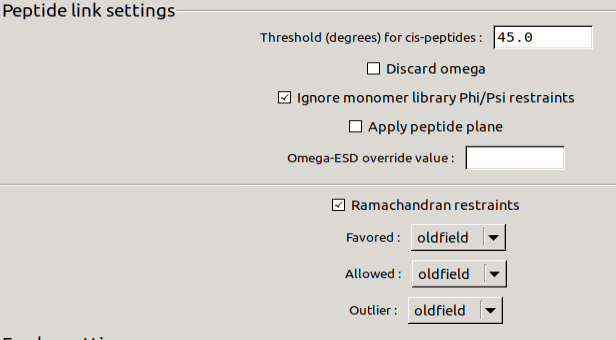

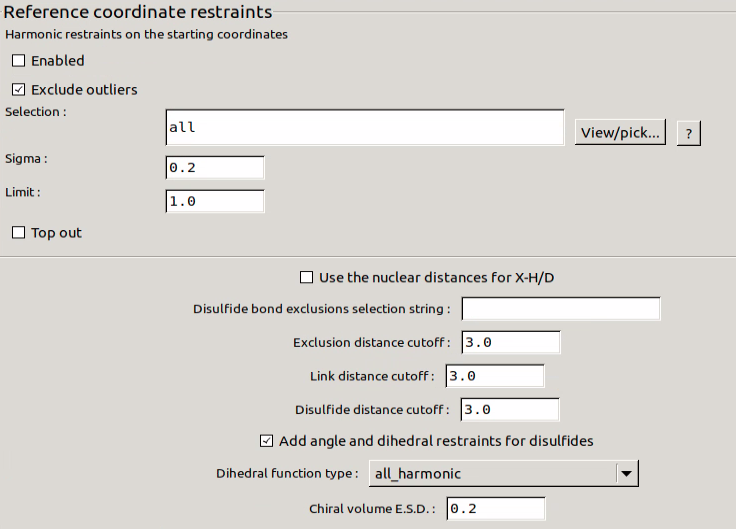

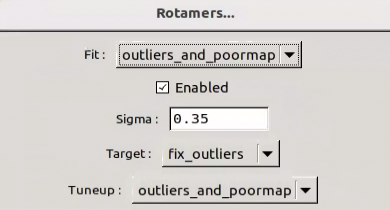

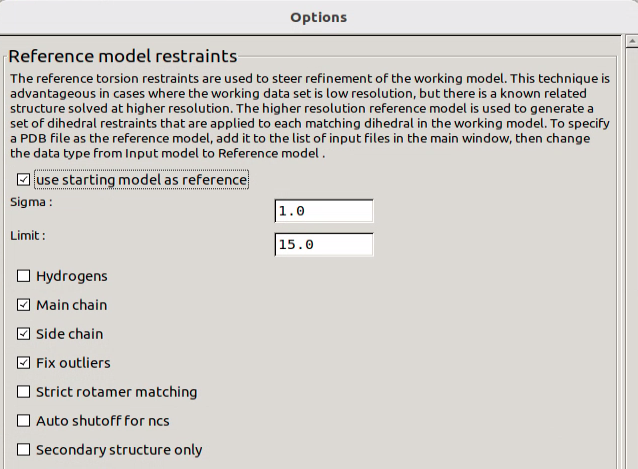

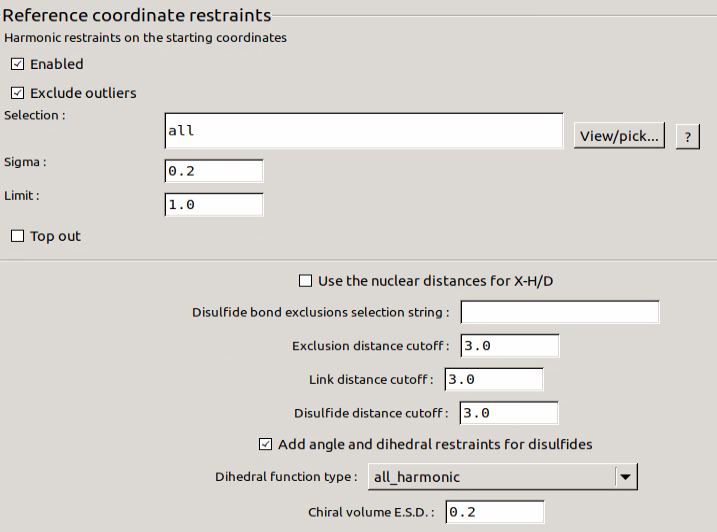

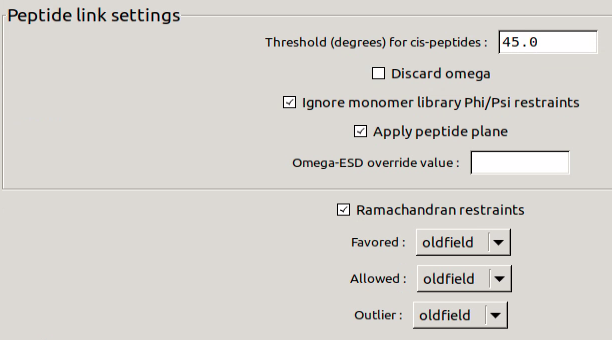

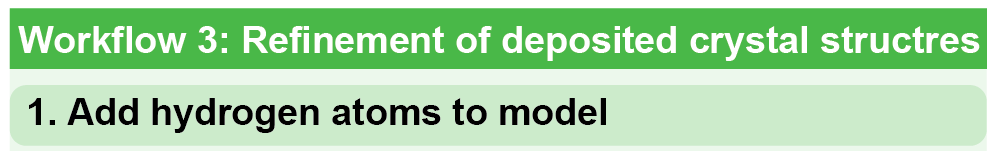

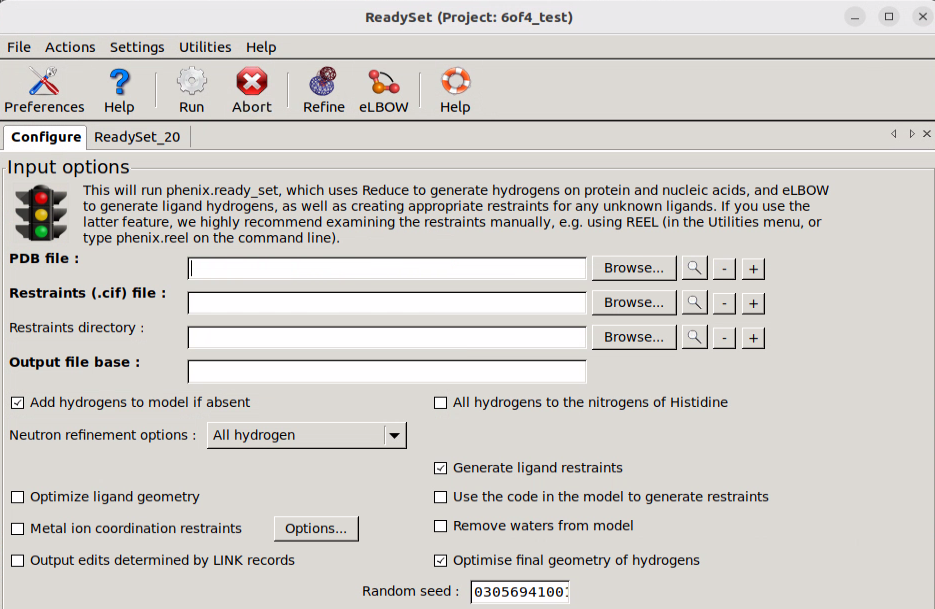

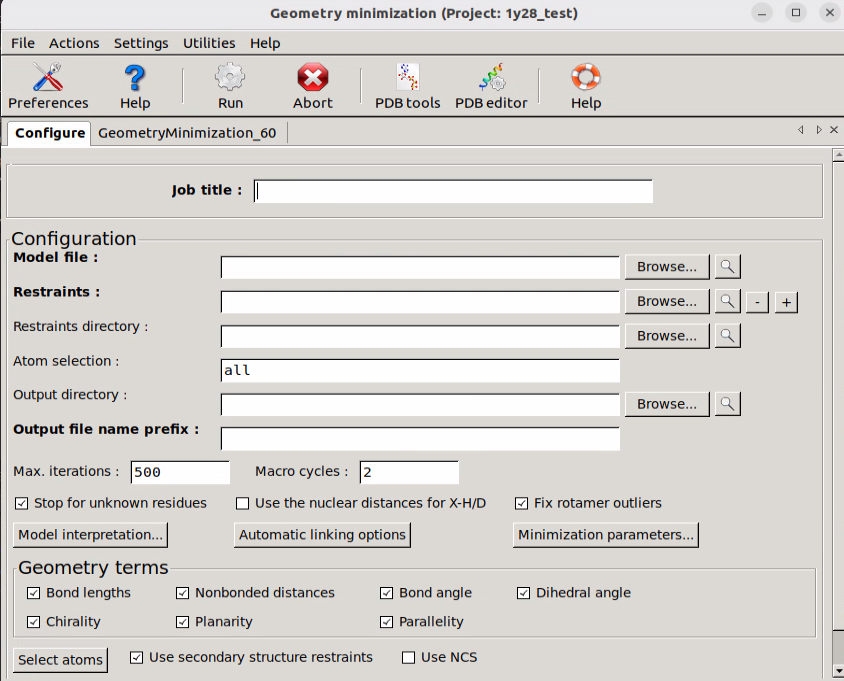

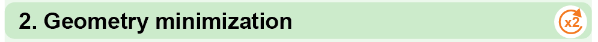

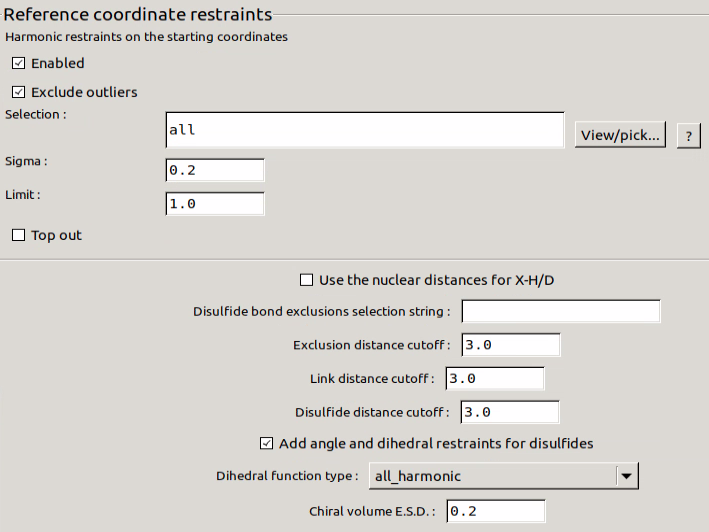

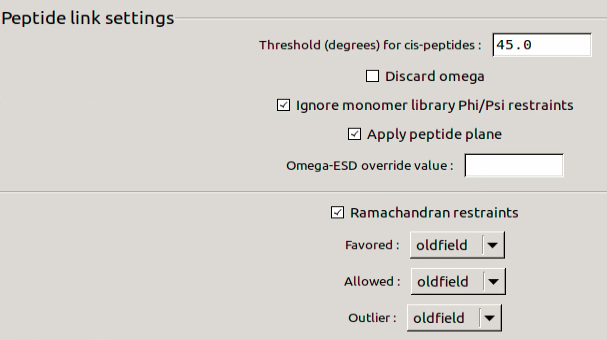

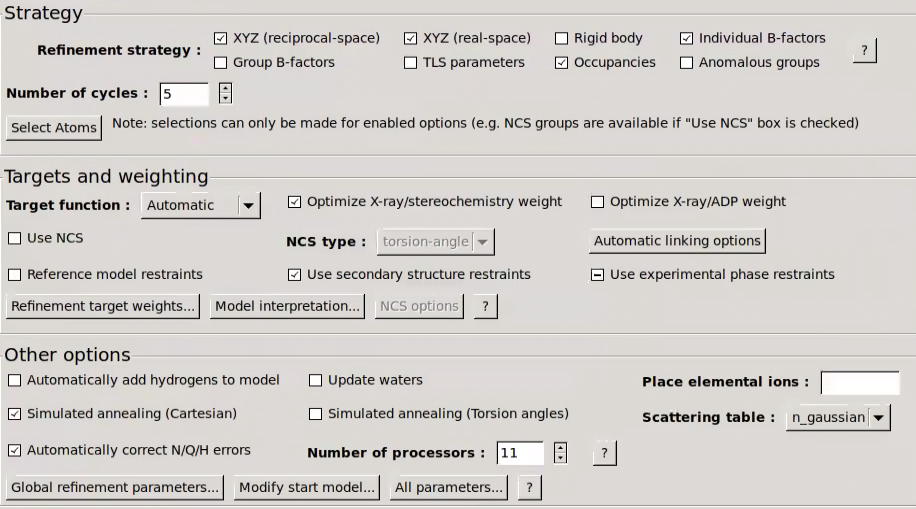

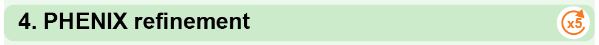

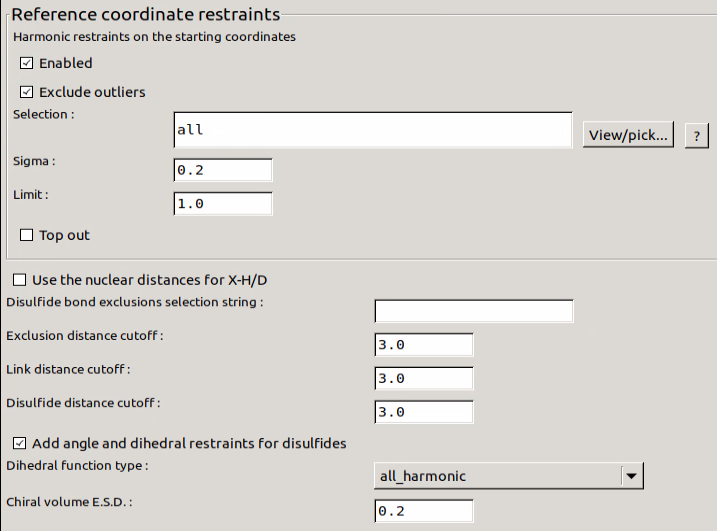

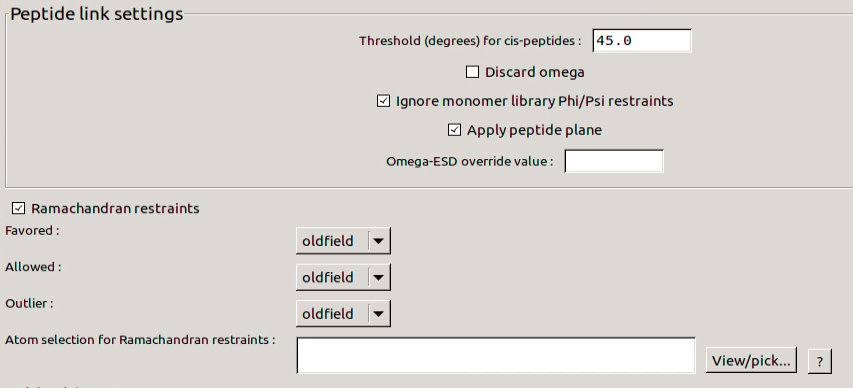

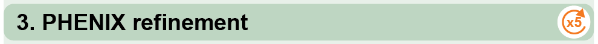

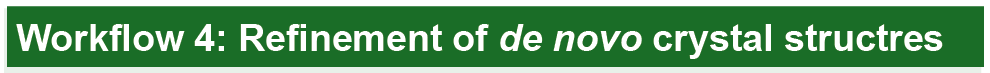

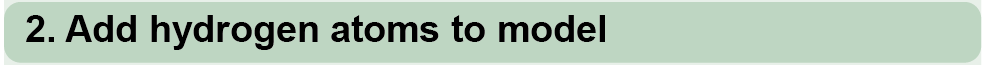

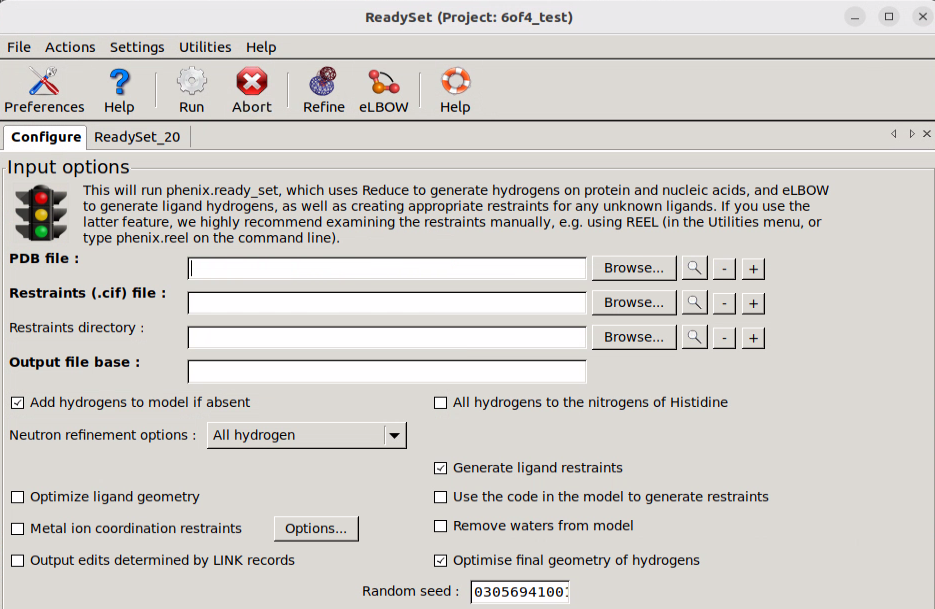

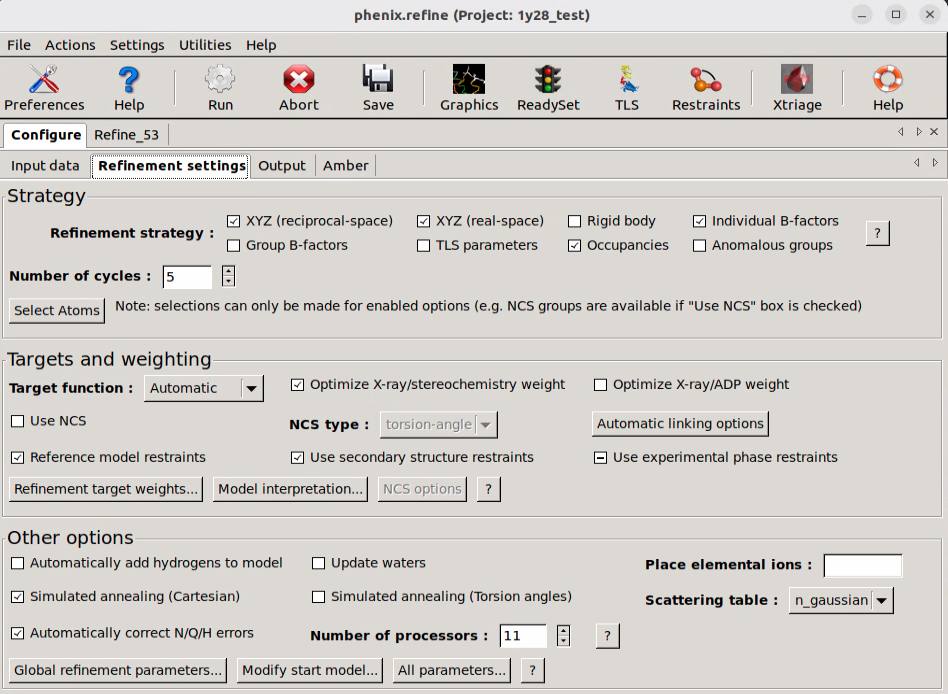

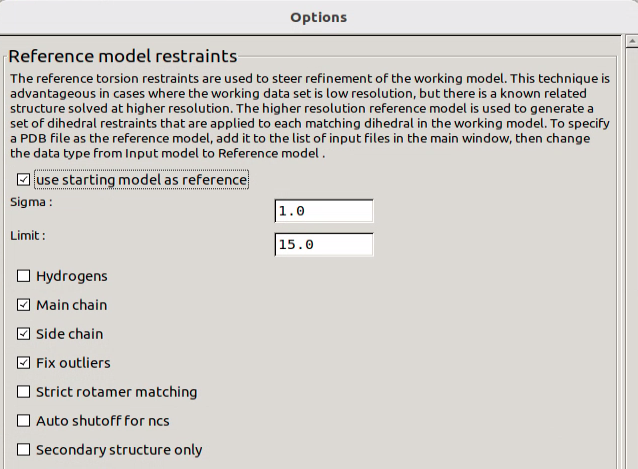

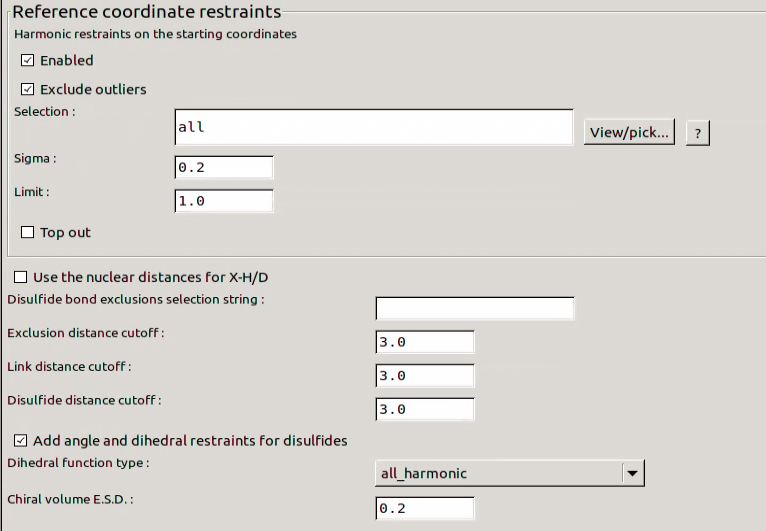

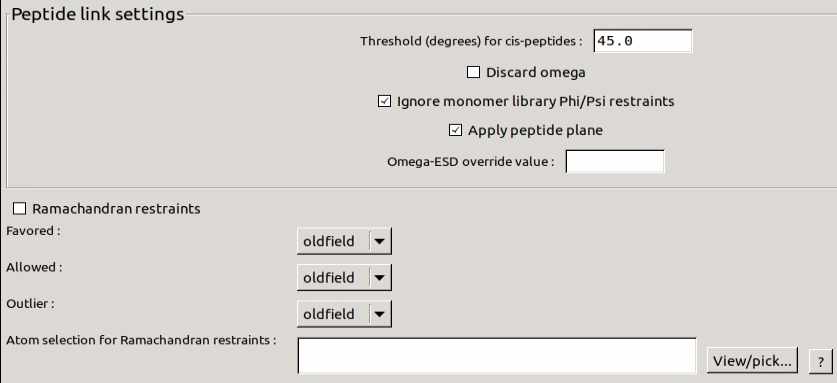

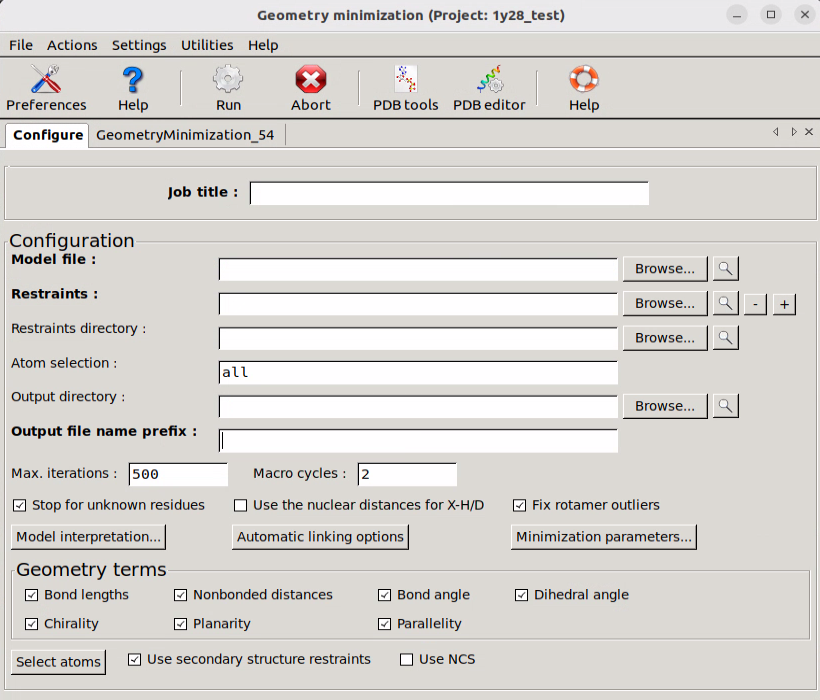

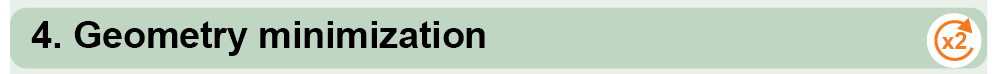

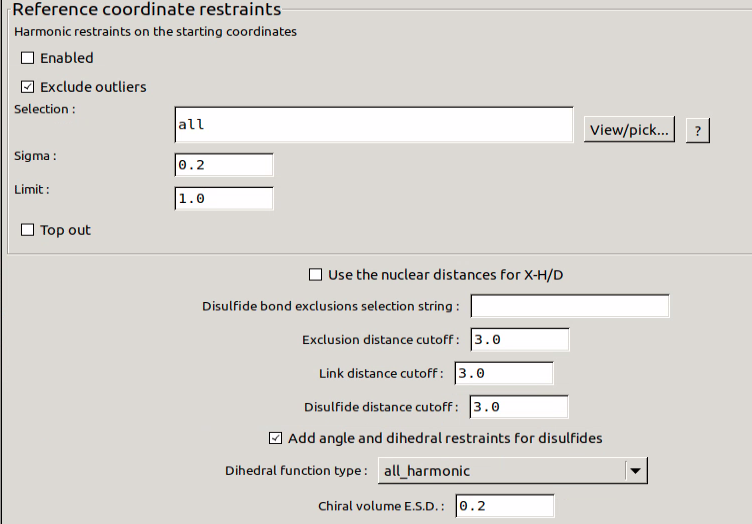

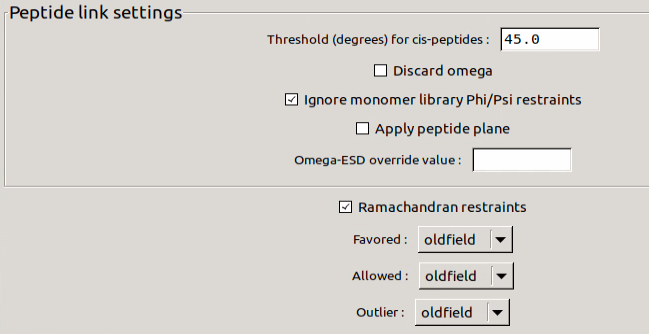

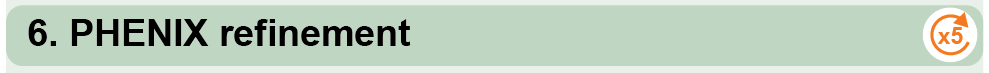

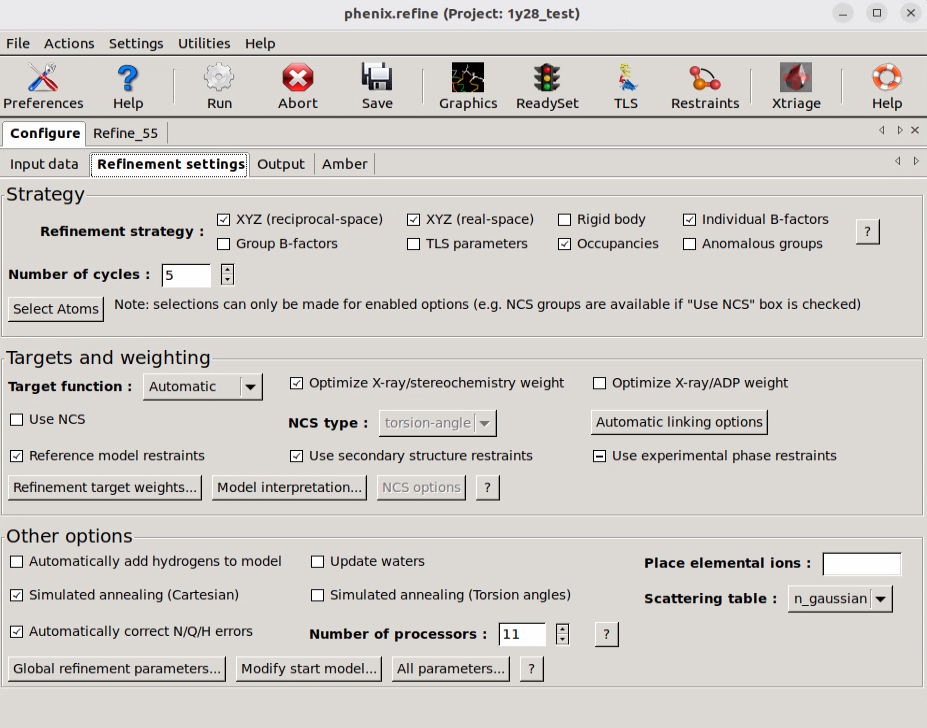

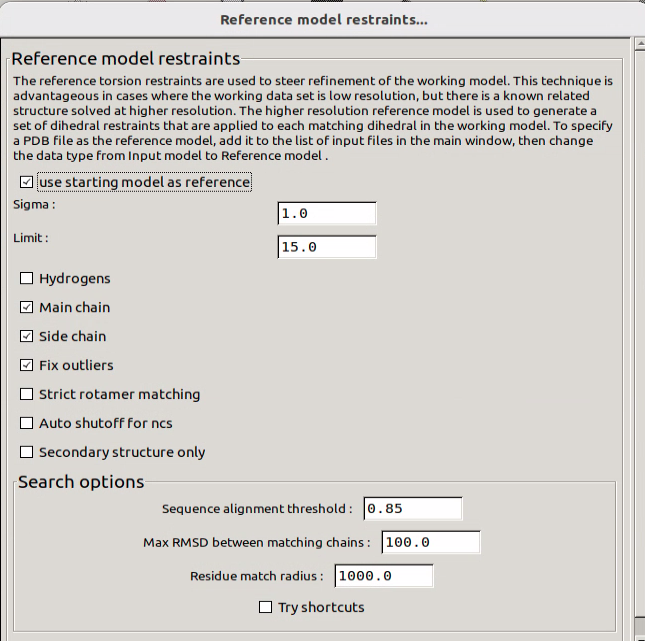

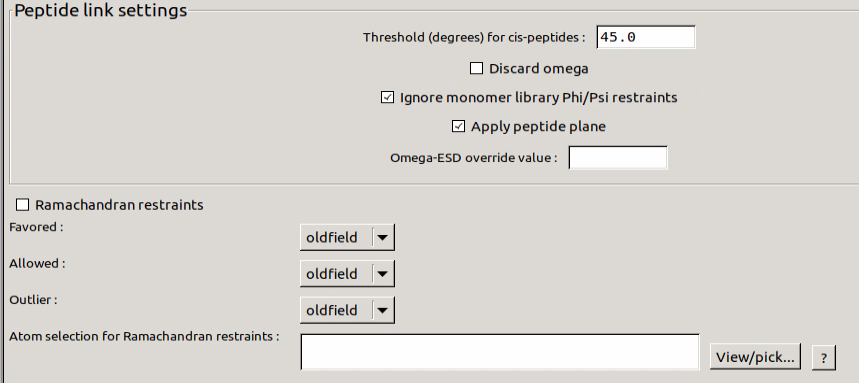

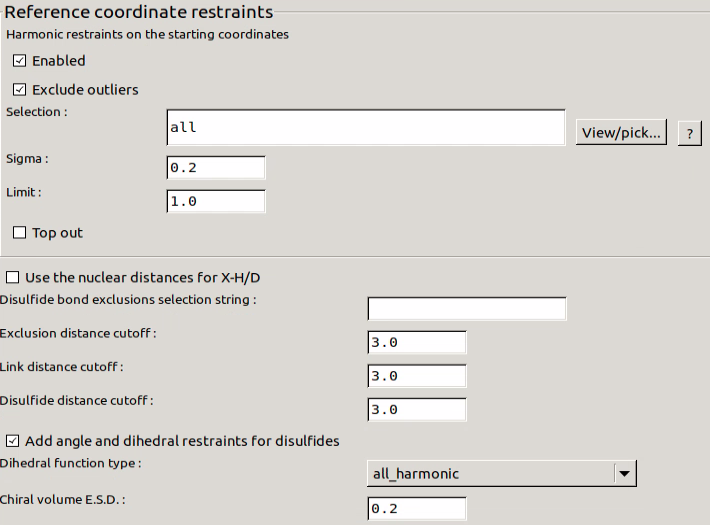
